# Generalizability of machine learning models for plant traits using hyperspectral reflectance data: The case of maize

**DOI:** 10.1101/2025.07.03.661070

**Authors:** Rudan Xu, John Ferguson, Johannes Kromdijk, Zoran Nikoloski

## Abstract

Hyperspectral reflectance provides rapid and precise phenotyping of plants in a non-destructive manner both in field and well-controlled settings. The resulting data have been used to devise machine learning (ML) models for paired measurements of different traits in diverse plants and crops. Yet, despite advances in using of hyperspectral data to reliably predict crop traits of interest, there are pressing issues concerning the training of ML models, the aggregation of data from crop field trials, and the generalizability of the models in different prediction settings. We collected hyperspectral reflectance data along with 25 anatomical, gas exchange, and chlorophyll fluorescence traits from 320 recombinant inbred lines of a maize Multi-Parent Advanced Generation Inter-Cross population grown across three consecutive seasons. We use these data to systematically: (1) compare the performance of representative ML models for different traits, including slow fluorescence kinetics whose predictability by hyperspectral data has not yet been investigated, (2) evaluate the ML model performance in prediction scenarios concerning unseen genotypes, unseen seasons, and the combination thereof, (3) investigate the effects of data aggregation of ML model performance. These problems are addressed in a rigorous nested cross-validation setting that provides a template for adequate assessment of performance of ML models for diverse crop traits considering the particularities of the experimental design.

**Significance statement:** We present a comprehensive evaluation of hyperspectral reflectance for predicting 25 physiological traits, including fluorescence kinetics, in maize across three seasons using rigorous nested cross-validation. By comparing ML models, prediction scenarios, and data aggregation strategies, our study reveals trait-specific limits of generalizability and offers a robust framework for deploying hyperspectral data in breeding applications.

## 1. Introduction

Efforts to develop climate-resilient crops go hand-in-hand with advances in high-throughput phenotyping technologies that can be effectively deployed in accelerated breeding (Kole et al., 2015; Langridge et al., 2021). Urgent innovations in high-throughput profiling are needed given that many breeding targets include molecular and biochemical traits that are challenging to measure in large breeding populations (Alseekh et al., 2021). Hyperspectral reflectance is one technology that provides rapid and precise phenotyping of plants in a non-destructive manner, allowing its deployment both in field and well-controlled settings (Grzybowski et al., 2021). Hyperspectral reflectance captures the intensity of light reflected from a plant across many specific wavelengths. The hyperspectral reflectance data (HSR) then consist of aggregated reflectance intensities at different light wavelengths. HSR data have been used to devise machine learning (ML) models for paired measurements of various molecular, biochemical, and physiological traits in diverse plants and crops (Kumar et al., 2001; Serbin et al., 2012; Yendrek et al., 2017; Silva-Perez et al., 2018; Cotrozzi et al., 2020; Feng et al., 2020; Furbank et al., 2021; Heckmann et al., 2017; Rehman et al., 2020; Wang et al., 2021; Yu et al., 2022; Kaur et al., 2024). Despite the wide application of HSR-based ML models, key methodological questions remain underexplored—particularly regarding model calibration, data aggregation strategies, relevant for breeding-related decisions, as well as generalizability across genotypes and growing environments.

With few exceptions, involving deep learning models (Rehman et al., 2020; Yu et al., 2022), most ML studies employ Partial Least Squares Regression (PLSR) as the standard model for trait prediction using HSR data as predictive features. A key hyperparameter in PLSR is the number of components (NoC), that must be carefully selected to avoid overfitting while ensuring model accuracy. The value for NoC is typically identified using a calibration data set. This calibration data set must not overlap with the validation dataset, used to assess model performance, to prevent data leakage. However, the strategy used for selecting NoC and model evaluation varies significantly across studies.

Two main approaches are commonly used to train and evaluate PLSR models–proposed by Burnett et al. (2021) and Filzmoser et al. (2009). The first approach, used, for instance, in Serbin et al. (2012, 2014), Meerdink et al. (2016), Ely et al. (2019) and Wang et al. (2021), involves random partitioning of the full dataset into a calibration set (typically 80%) and a validation set (20%). Within the calibration set, 70% of the samples are randomly selected to train the model using varying values of NoC, while the remaining 30% of the calibration set are used to assess model performance of each NoC. This procedure is typically repeated 1000 times to ensure robust estimation. The optimal NoC value is chosen by minimizing the predicted residual error sum of squares (PRESS) statistics (Chen et al., 2004) collected from the repeated sub-sampling. The second approach, known as repeated double cross-validation, employs two nested cross-validations, and has been utilized in studies such as Dechant et al. (2017), Yan et al. (2021) and Ji et al. (2024).

In this case, an outer loop repeatedly partitions the dataset into calibration and validation folds (e.g., 5-fold CV repeated 100 times). An inner loop performs k-fold CV within each calibration set to identify the optimal NoC using the standard error method (Tibshirani & Friedman, 2001). This method prevents the selection of excessively large NoC that might reduce the mean squared error (MSE), while increasing the risk of overfitting.

The assessment of the generalizability of the resulting models to unseen samples has been approached in different ways. For instance, Burnett et al. (2021) assessed the uncertainty of PLSR models using jackknife (leave-on-out) of the calibration data set, given that a single final validation set was available. In contrast, Filzmoser et al. (2009) proposed repeated cross-validation sets, allowing thorough assessment of both prediction accuracy and performance variability. It is then conceivable that these differences can have impact on the findings and their comparability between different studies.

Here, we collected HSR data and paired measurements of 25 traits from 320 recombinant inbred lines (RILs) of a maize Multi-Parent Advanced Generation Inter-Cross (MAGIC) population grown across three consecutive seasons. Using this dataset, we addressed the following key questions:

1. How can one select the optimal NoC in PLSR in an accurate and computationally efficient way? To address this question, we systematically compared different inner-loop structures and selection criteria using repeated cross-validation to assess their impact on model performance and uncertainty.
2. How do ML models perform under increasingly challenging scenarios of generalization? To this end, we evaluated three levels of model generalization: (i) random splits of training and validation data; (ii) prediction of unseen genotypes within the same season; and (iii) prediction of traits from unseen seasons (Figure 1a). We note that these scenarios of model generalizability are particularly relevant for making decisions based on the ML models in breeding.
3. How do data aggregation and season influence model performance? In this respect, we investigated the impact of averaging raw trait and reflectance measurements at the plot or genotype level. In addition, we investigated how aggregation affects the model generalization scenarios, mentioned above. To this end, we made use of the three replicates of the genotypes per plot, to compare models that are based on HSR and traits data aggregated by averaging over plots or averaging over genotypes.

**Figure 1.**
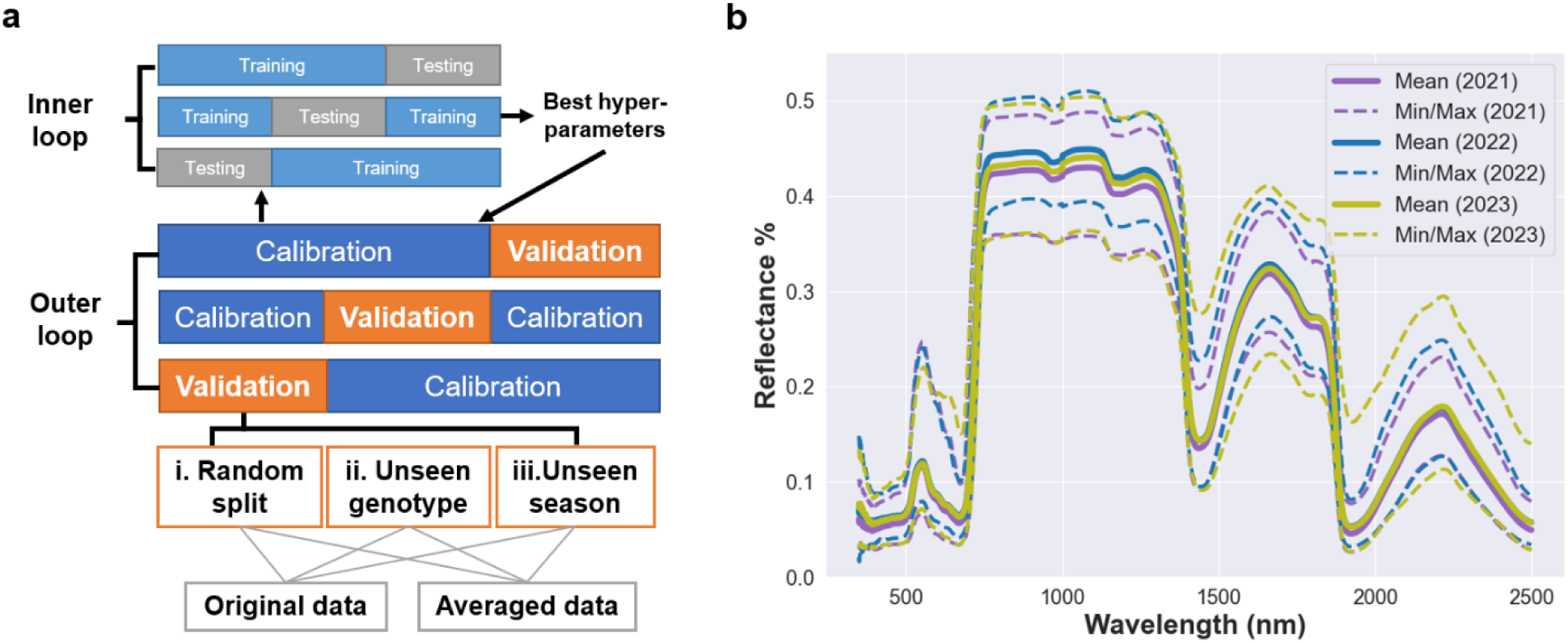
Illustration of the machine learning framework for prediction of physiological traits using HSR data. (a) Repeated nested cross-validation (CV). Outer CV loop is used for splitting the data set into a calibration set and a validation set; the latter is used to assess model performance under different prediction scenarios, namely: random unseen samples, unseen genotypes, and unseen seasons. Within th calibration sets, inner CV loop is used to tune hyperparameters. (b) Summary of the ranges of HSR data measured on 320 lines from a maize MAGIC population grown in field experiments in three consecutive seasons, i.e., 2021, 2022 and 2023. The mean HSR values over the genotypes in each season are depicted by a bold line, while the minimum and maximum values are plotted with dashed lines across th measured wavelengths.

By addressing these questions, our study provides a robust experimental and analytical framework for assessing model performance on HSR data from field trials, with direct implications for predictive phenotyping across genotypes, traits, and environments.

## 2. Materials and methods

### 2.1 Leaf physiological traits and reflectance measurements

Field trials incorporating the maize MAGIC population (Dell’Acqua et al., 2015) were performed in 2021, 2022, and 2023 at the National Institute of Agricultural Botany (NIAB, Cambridge, UK). In each year, 320 recombinant inbred lines (RILs) were grown in a twice-replicated alpha lattice design. The experimental design has previously been described in full detail (Ferguson et al., 2025). Some RILs were specific to certain years, but 305 RILs were grown and phenotyped in all three years.

All phenotyping was performed on the leaf subtending the ear, which was excised at dawn and returned to the laboratory for phenotyping, as described previously (Ferguson et al., 2025). We have previously shown that this approach generated data comparable to measuring leaves still intact to the plant (Ferguson et al., 2023).

Initially, light-saturated photosynthesis (*A*) was determined on all leaves using LI-6400 infra-red gas analyzers equipped with 6400-40 leaf chamber fluorometer LED light sources (LI-COR Inc., Lincoln, NE). Conditions within the leaf chambers were as follows: 1800 µmol m-2 s-1 photosynthetically active radiation (PAR), 25°C block temperature, 400 µmol s-1 air flow, 65% relative humidity (RH), 400 µmol mol-1 reference CO_2_ concentration. A randomly selected subset of RILs (78 in 2021, 88 in 2022, 91 in 2023) were used for measuring the response of photosynthesis to changes in the intracellular concentration of CO_2_ (*A-Ci* curve) using LI-6800 infra-red gas analysers equipped with standard 6cm2 leaf chambers (LI-COR Inc., Lincoln, Nebraska). *A-Ci* response curves were collected exactly as described previously (Ferguson et al., 2024). Using the *A-Ci* data, estimates of the maximum rate of carboxylation by phosphoenolpyruvate carboxylase (PEPC; V_pmax_) and the asymptote of the *A-Ci* curve (V_max_) were modeled following von Caemmerer (Von Caemmerer, 2000).

Leaf hyperspectral reflectance was measured on the same portion of the leaf where gas exchange was performed using an ASD FieldSpec 4 Standard-Res Spectroradiometer equipped with a leaf-clip (Malvern Panalytical, Malvern, UK). The light source of the FieldSpec was allowed to warm up for 45 minutes prior to measurements. For each leaf, three technical replicates were performed and the reflectance at each wavelength was averaged before subsequent data analyses.

Following hyperspectral reflectance, a section of leaf tissue (∼2×4 cm) was excised from the previously measured leaf area for measurements of chlorophyll fluorescence to determine the quantum efficiency of photosystem II (Fv/Fm) as well as the response of non-photochemical quenching (NPQ) and photosystem II (PSII) operating efficiency (ΦPSII) to an actinic light being switched on and off. These measurements are as described previously (Ferguson et al., 2025). Traits describing NPQ induction and relaxations, as well as ΦPSII recovery responses were obtained from exponential models. The 2021 and 2022 chlorophyll fluorescence data have previously been reported (Ferguson et al., 2025).

### 2.2 Data processing

For each genotype, measurements of different traits were collected from two plots, with three replicates each. To assess differences in model performance, raw data were averaged by plot or genotype. Wavelengths from 400 nm to 2400 nm were used in training models to predict each of the traits, except for nitrogen percentage and ratio of nitrogen to carbon percentage. For these traits, wavelengths from 1500 nm to 2400 nm were used, as in Yendrek et al. (2017). To account for the variability of trait response across seasons, particularly for SLA, before combining different seasons as training data, the trait values from each year were centered and scaled.

### 2.3 Machine learning models

Supervised machine learning (ML) was used to develop models to predict measured traits based on HSR data. Once trained, these models can predict traits for which only reflectance data are available. Given that the traits are continuous variables, regression-based ML models were used. The predictions were performed by using ‘pls’ (Mevik & Wehrens, 2023) and ‘caret’ package (Kuhn & Max, 2008), implemented in R software environment.

Partial least-squares regression (PLSR) is a state-of-art ML model commonly used in modeling plant traits based on HSR data as predictors (Geladi & Kowalski, 1986; Wold et al., 2001). HSR typically contain more predictors (i.e., reflectance at different wavelengths) than samples, and these predictors are often highly correlated, leading to a multicolinearity problem. PLSR addresses these challenges by projecting the response variable onto uncorrelated components, which are linear combinations of predictors. This approach allows the response to be expressed in terms of the original predictors by considering the response during component extraction. The number of components is a crucial hyperparameter in PLSR and is typically tuned using different procedures, as described above. PLSR has been successfully used to predict traits such as V_max_, J_max_ and %N across different species (Yendrek et al., 2017; Meacham-Hensold et al., 2019).

For fairness of comparison to the widely used PLSR models based on HSR data, we considered linear kernel SVR to avoid nonlinearity of other kernels as explanation for differences in performance. As a result, the SVR also has one hyperparameter, *C*, referred to as the regularization hyperparameter. In this study, we implemented a nested CV workflow to tune the hyperparameter and validate the model using unseen data under different scenarios.

### 2.4 Different prediction scenarios

This study evaluated the utility of HSR data for predicting physiological traits under three distinct prediction scenarios, each designed to assess model generalizability under varying levels of data partitioning. Model performance was assessed using the outer loop of a repeated CV framework, while hyperparameter tuning was conducted within the inner loop, as illustrated in Figure 1a. The scenarios are described in the following:

Scenario i: Random sample prediction. In this scenario, model performance was evaluated using randomly partitioned data sets. This approach enables systematic comparison of model accuracy across different data aggregation strategies and seasons. The outer loop consists of 20 repetitions of random 5-fold CV, allowing the collection of mean performance from five folds across repetitions. Hyperparameter optimization within the inner loop was conducted using either repeated CV, permutation-based, or single CV approaches.

Scenario ii: Unseen genotype prediction. A 5-fold CV scheme was repeated 20 times, ensuring genotype exclusivity between folds in both outer and inner loops to avoid data leakage. This set-up allowed for the assessment of the model’s performance across genotypes.

Scenario iii: Unseen season prediction. In this scenario, the goal was to predict traits for a new, unseen season using data from other seasons in model training. Here, entire data sets from one or two seasons were used for model calibration, and model performance was evaluated on data from the held-out season. Within the calibration set, the inner loop employed random splits for hyperparameter tuning. Additionally, a more stringent variation of this scenario was implemented, in which the model was trained on data from a single season and was then applied to unseen genotypes in an unseen season. This was achieved by withholding 1/5 of the data of the genotypes from the training season and evaluating their predictions in a separate season in the framework of 20 repetitions of 5-fold CV. Here, too, genotype exclusion was strictly enforced during inner loop splitting to avoid data leakage.

Model performance was quantified by using coefficient of determination (R^2^) for scenarios i and ii and squared Pearson correlation (r^2^) for scenario iii. We used squared Pearson correlation (r^2^) to assess the trend of predicted values rather than their agreement on measured values in the most challenging scenario iii.

## 3. Results

### 3.1 Phenotyping a maize MAGIC population gathers large-scale data suitable for testing the generalizability of ML models

To address our questions, we gathered leaf HSR data obtained from a maize MAGIC population grown in three consecutive field seasons (i.e., 2021, 2022 and 2023) (Figure 1b, Methods). The average HSR profiles from three seasons over all samples were consistent across the measured wavelength region of 700 to 3500 nm. In the visible region (350–700 nm), both the minimum and maximum HSR values over all samples were similar across the seasons. However, by calculating the correlation of averaged HSR per genotype, the mean correlation based on the data from this region ranged from 0.27 to 0.36 between three seasons (Table S1). In the near-infrared region (700–1300 nm), where HSR reaches the high plateau, the minimum values measured in 2022 were significantly higher than those measured in 2021 and 2023, while the maximum values in 2021 were lower than those in 2022 and 2023 (Figure 1b). The average correlation for this region across genotypes between seasons 2022 and 2023 was 0.5, which was lower than the correlation of 0.57 observed between the 2021 and 2022 seasons and the 0.53 observed between the 2021 and 2023 season. The shortwave infrared region (1300–2500 nm) also showed differences in the range of HSR values across the three seasons, with averaged correlation of 0.50, 0.44 and 0.49 for the pairs of three seasons (Table S1). Moreover, Mantel tests were performed to assess if the covariance matrix of HSR data (across genotypes) in one season were correlated with the covariance matrix from another season. We found that seasons 2021 and 2022 showed a correlation of 0.89, seasons 2021 and 2023 had a correlation of 0.88 and seasons 2022 and 2023 showed a correlation of 0.93. These values indicated differences in the covariance structure of the HSR data that may affect transferability of the models, particularly involving training / testing with data from season 2021 that differs the most from data from 2022 and 2023.

In addition, 25 traits were measured or estimated during these seasons, with their distribution depicted in Figures S1, S2 and S3. These traits were grouped into three categories, with different number of samples in each category (Table S2). We measured six anatomical traits, including: specific leaf area, relative C content (%C), relative N content (%N), carbon to nitrogen ratio (C:N) as well as the stable carbon and nitrogen isotope ratios ( δ^13^C and δ^15^N). Seven gas exchange traits were estimated from the Farquhar-von Caemmerer-Berry (FvCB) model (Von Caemmerer, 2000) using *A-Ci* response curves, namely: stomatal limitation, photosynthetic rate, stomatal conductance, and intrinsic water use efficiency (iWUE) at saturating light, maximum rate of carboxylation by phosphoenolpyruvate carboxylase (V_pmax_), asymptote of the *A-Ci* curve (V_max_). Moreover, we considered 13 chlorophyll fluorescence traits, related to quantum efficiency of PSII (Fv/Fm), response of non-photochemical quenching (NPQ) and photosystem II operating efficiency (ΦPSII).

We observed that δ^15^N, stomatal limitation, and NPQ induction rate exhibited the highest average coefficient of variation (CVs) within season (CVs>45%), while δ^13^C, Fv/Fm and final ΦPSII showed CVs below 5%. We also observed that V_pmax_ had higher variability compared to V_max_, with average CVs of 39% and 24%, respectively. To assess the consistency of the measurements across seasons, we determined the Spearman correlation between seasons for each trait, averaged per genotype (Table S3). We found that, %C, δ^15^N and iWUE, on average, had the lowest correlation between seasons, with mean correlation values of 0.02, 0.21 and 0.24, respectively. In contrast, fluorescence traits such as NPQ relaxation amplitude, ΦPSII recovery offset and ΦPSII induction amplitude showed the highest correlation between seasons, with values of 0.60, 0.62 and 0.64. Interestingly, SLA and C:N had correlation of 0.43 and 0.53 while V_pmax_ and V_max_ averaged correlations of 0.32 and 0.47. These results suggested variability in both HSR and traits data between seasons, indicating presence of genotype-by-environmental interaction, that may severely limit the transferability of models between seasons.

### 3.2 PLSR model sensitivity to NoC estimation

In this section, we used the entire dataset from season 2021 as calibration data to evaluate three approaches for NoC estimation with different data partitioning strategies (CV or sub-sampling) and selection criteria (MSE-based or PRESS-based), followed by an evaluation of PLSR model performance to variations in NoC. For each trait, a fixed design of 20 repetitions of 5-fold CV was implemented to systematically assess model performance across a range of NoC values from 1 to 40. Model calibration performance across all 20 × 5 test folds was quantified using MSE and PRESS statistics. With the MSE-based selection criteria (Tibshirani & Friedman, 2001), the optimal NoC, NoC_MSE(opt)_, was selected for each repetition, and the most frequently chosen value across 20 repetitions was defined as the final estimated NoC: *NoC_MSE_*(Filzmoser et al., 2009). The distribution of NoC_MSE(opt)_ values thus captures variability of NoC selection under different data partitions and allows assessment of its impact on model performance. For comparison, a second approach using 100 random 80/20 training-testing partitions was employed to estimate NoC based on the PRESS statistic (Chen et al., 2004).

For this analysis, the full hyperspectral reflectance (HSR) ranged from 400 to 2400 nm collected in the 2021 season was used. Exceptions were made for %N and C/N, for which only the 1500–2400 nm range was used, similar to previous studies (Serbin et al., 2014; Yendrek et al., 2017).

Overall, NoC estimates were largely consistent between these two methods. Difference between *NoC_MSE_* and *NoC_PRESS_* were observed in twelve traits, with the largest discrepancies (Δ=4) seen for SLA, NPQ induction slope, and Fv/Fm. Median R^2^ differed by less than 0.03 across all traits, suggesting strong agreement between the two selection strategies. Notably the PRESS-based method tended to favor slightly smaller NoC values for some traits (Tables S4 and S5).

Using the MSE-based method, the standard deviation of NoC_MSE(opt)_ across 20 repetitions ranged from 0 to 3 for 22 traits, indicating stable NoC estimation (Table S4). In contrast, δ^13^C and stomatal conductance (gsw) showed slightly higher standard deviation of 4, with corresponding median R^2^ values of 0.12 and 0.23 respectively. Notably, δ^15^N and NPQ induction rate exhibited the highest the standard deviation (7) and had near-zero median R^2^, suggesting unstable NoC selection likely due to the lack of predictive signal in the HSR data.

To compare model calibration performance, median R^2^ was computed across all 20 × 5 test folds using a single consensus NoC (*NoC_MSE_* ), while median R^2^ was calculated by applying the individually selected NoC_MSE(opt)_ for each repetition. The results showed that twelve traits yielded slightly higher median R^2^ values using NoC_MSE(opt)_, five traits performed better with NoC_MSE_, and the remainder showed no difference. Additionally, the standard deviations of R and R^2^ were nearly identical, suggesting that most of the performance variability across iterations originated from differences in data partitioning rather than variation in NoC_MSE(opt)_. Although NoC estimates may vary slightly across repetitions, such variability had small effect on overall model calibration performance, which generally plateaued within a stable NoC range (Figure 2).

**Figure 2.**
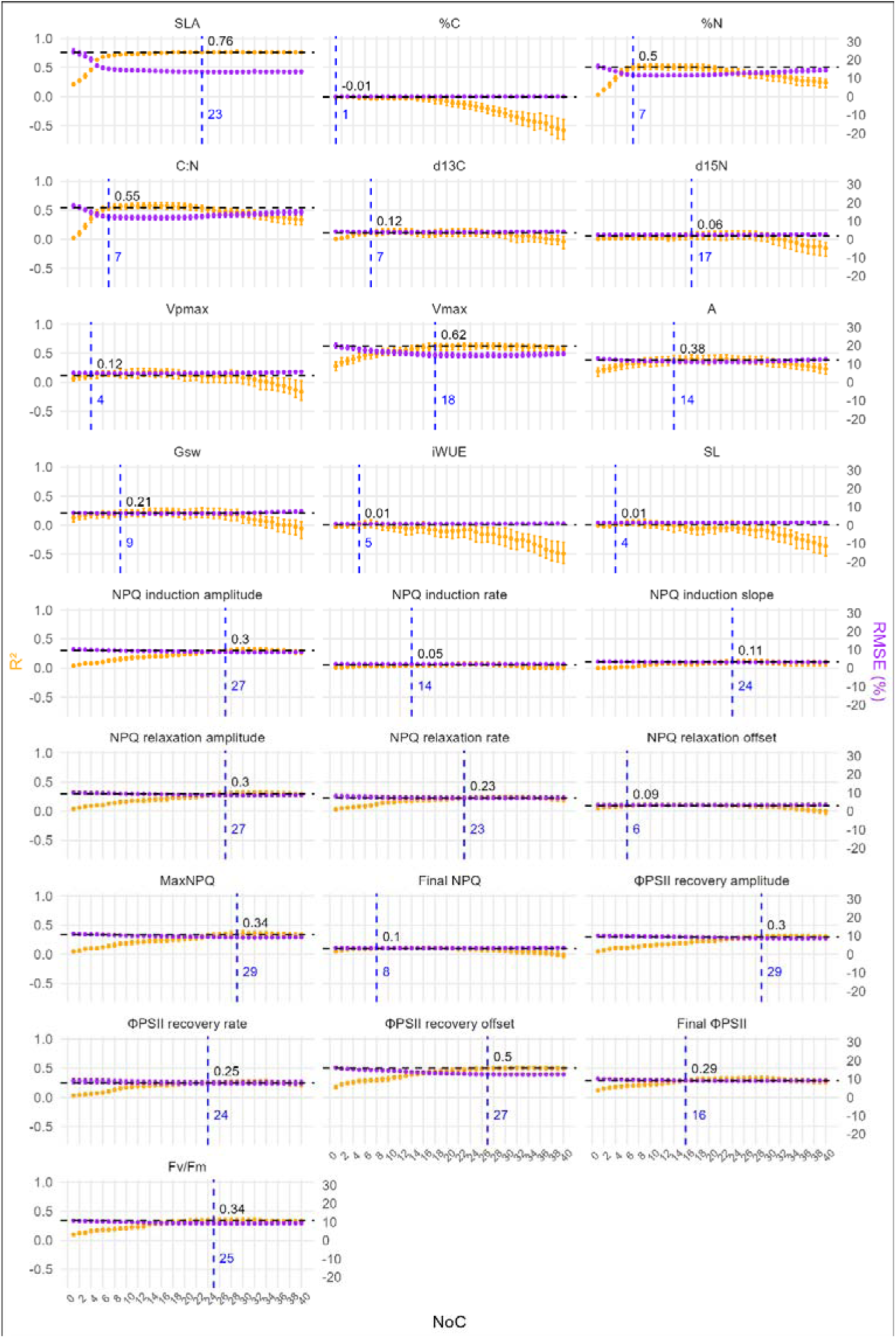
Sensitivity of model performance to the number of component (NoC) for PLSR model across 27 physiological traits measured in season 2021. For each trait, model performance was evaluated using 20 repetitions of 5-fold cross-validation across varying NoC values. Performance metrics included the coefficient of determination (R^2^, orange, left y-axis) and RMSE% (purple, right y-axis). Dots indicate the median score across the 100 test folds, with error bars representing the interquartile range (25th to 75th percentile). Vertical dashed lines mark the *NOC_MSE_*, while horizontal dashed lines denote the corresponding median R^2^.

We classified the 25 traits into four categories based on the shape of their R^2^ response across increasing NoC (Figure 2):

1. Traits with stable and robust performance across NoC. These traits exhibited increasing R^2^ and decreasing RMSE% with increasing NoC, followed by a performance plateau where model accuracy stabilized and variability across CV splits was small (Figure 2 and Table S4). SLA showed the best performance overall (median R^2^=0.76, *NoC_MSE_* =23). Other traits that could be predicted well included V_max_ (median R^2^=0.62, *NoC_MSE_* =18) and several chlorophyll fluorescence traits such as ΦPSII recovery offset, ΦPSII induction amplitude, maximum NPQ, NPQ induction amplitude, NPQ relaxation amplitude and Fv/Fm, which achieved median R^2^ between 0.30 and 0.50, with estimated *NoC_MSE_* exceeding 20. While some traits like NPQ relaxation rate, NPQ induction slope, final ΦPSII and ΦPSII induction rate showed a median R^2^ ranged between 0.11 to 0.29. These traits maintained robust model performance even at high NoC values, with small variation across repetitions.

2. Traits with moderate predictability but overfitting beyond optimal NoC. For this group, R^2^ improved initially but declined beyond the optimal NoC region, indicating overfitting. This pattern was evident in traits such as %N, C/N, δ^13^C, A, gsw, which generally showed relatively higher variability across CV splits. Their median R^2^ at *NoC_MSE_* ranged between 0.12 and 0.56, suggesting that the model captured a meaningful predictive signal, despite the eventual decline in performance due to overfitting at higher levels of model complexity.

3. Traits with low predictability and overfitting beyond optimal NoC. In contrast, traits including %C, δ^15^N, V_pmax_, Stomatal limitation and iWUE exhibited low R^2^ (< 0.15) at low NoC and decreasing R^2^ at higher NoC, accompanied by large variability across CV splits. These results suggest that the models could detect weak predictive signals for these traits, that, however, do not generalize well.

4. Traits with flat and low R^2^ curves across NoC. These traits exhibited consistently low R^2^ values across the entire range of NoC, with minimal variation across CV splits. This flat trajectory, observed for traits such as NPQ induction rate and final NPQ, suggests that the model was unable to capture any meaningful relationship between the predictors and the response variable. Consequently, predictions were limited to the mean response, and increasing model complexity did not lead to performance deterioration.

Altogether, the estimated *NoC_MSE_* yielded the smallest RMSE and highest R while preventing overfitting for all traits (Figure 2). We also compared the model calibration using complete HSR versus a sub-sampled version in which every fifth wavelength was retained. The resulting median R^2^ and optimal NoC estimation were highly similar between the two approaches, indicating that wavelength sub-sampling offers an efficient and effective form of dimension reduction without compromising predictive accuracy (Table S6). Consequently, the sub-sampled HSR data were used for all downstream analyses.

In this analysis, the complete 2021 dataset was used to estimate the optimal number of components (NoC) and evaluate model calibration. Three calibration strategies were applied: single CV to determine NoC_MSE(opt)_, repeated CV for *NoC_MSE_*, and permutation-based splitting for *NoC_PRESS_* . These calibrated models can then be applied to predict data from independent seasons. However, to rigorously assess the model’s generalizability within the same season, a nested CV framework is required to ensure a proper separation between NoC estimation and model performance evaluation.

### 3.3 Generalizability of PLSR and SVR using different pre-processing techniques

To evaluate PLSR model generalizability within a season, we employed a 5-fold CV repeated 20 times as the outer evaluation loop, following the framework proposed by Filzmoser et al. (2009). In each iteration, one fold served as the validation set while the remaining data were used for calibration. Within the calibration set, two inner-loop strategies were compared for estimating the number of components (NoC): 10 times repeated 3-fold CV to derive the final *NoC_MSE_*, and a single CV iteration to obtain NoC_MSE(opt)_.

As anticipated, NoC_MSE(opt)_ exhibited greater variability across iterations than *NoC_MSE_*, but this did not significantly impact the final validation R^2^ (Figure S4). Strong predictive performance (median R^2^ > 0.5) was observed for traits such as SLA, %N, C:N ratio, and V_max_. Moderate performance (0.3 < median R^2^ < 0.5) was achieved for A, NPQ induction and relaxation amplitudes, ΦPSII recovery amplitude and offset, final ΦPSII, Fv/Fm, and maximum NPQ.

The dataset from season 2021 contained up to six HSR-trait measurement pairs per recombinant inbred line (RIL): three from each replicate (plot) of the alpha-lattice design. These raw data were aggregated either by plot or by genotype. To explore the effect of data aggregation as a pre-processing method on model performance, we applied PLSR using sub-sampled HSR data with repeated CV for NoC estimation. Notably, genotype-averaged data improved model accuracy for traits such as SLA, %N, and C:N ratio, with median R^2^ increasing from 0.75, 0.50, and 0.56 (raw) to 0.81, 0.74, and 0.74, respectively (Figure S5). Most photosynthesis traits performed better using raw data, except for V_max_, which improved by plot-level averaging (median R^2^=0.60). Whilst most fluorescence traits performed best with raw data, ΦPSII recovery offset, final ΦPSII and Fv/Fm showed improved predictive accuracy when using plot-averaged data, with median R^2^ increasing from 0.49, 0.30 and 0.32 to 0.5, 0.38 and 0.42, respectively. Aggregation also influenced NoC estimation: genotype averaging typically reduced estimated NoC values, while still improving prediction accuracy for traits such as %N and C:N ratio (Figure S5).

Availability of data from experiments conducted across three consecutive seasons enabled us to evaluate whether the observed effects of data aggregation were consistent for the other two seasons. For SLA, genotype-averaged data consistently outperformed raw data across all three seasons, confirming the robustness of this aggregation strategy (Figure S6). Due to the absence of raw biochemical traits measurements for 2022 and 2023, only plot- and genotype-level aggregations could be evaluated. Notably, genotype-level averaging led to the most substantial improvements for %N and the C:N ratio, increasing their median R^2^ values from 0.64–0.68 and 0.71–0.73 (with plot-level data) to 0.72–0.74 and 0.78–0.79, respectively. In contrast, photosynthesis and most of chlorophyll fluorescence traits did not benefit from data aggregation. For these traits, both plot- and genotype-level averaging led to reduced predictive accuracy compared to models trained on raw observations. This suggests that preserving fine-scale variability may be crucial for capturing the spectral signals associated with physiological processes. An exception to this trend was observed for ΦPSII recovery offset, final ΦPSII and Fv/Fm, which consistently showed improved predictability with plot-averaged data across all three seasons.

Traits such as %C, δ^15^N, iWUE, stomatal limitation, NPQ induction rate and slope, NPQ relaxation offset, and final NPQ consistently showed poor predictability (maximum median R^2^ < 0.15 across seasons and aggregation methods) and were therefore excluded from further analysis.

To benchmark our findings against another linear method, we applied support vector regression (SVR) with a linear kernel using the same repeated CV framework. SVR yielded comparable results to PLSR calibrated with single CV, and outperformed it slightly for traits including SLA, δ^13^C, V_pmax_, A, and gsw measured in season 2021 (Figure 3). The consistency in aggregation effects across both PLSR and SVR suggests that the improvements observed for SLA, %N, C:N ratio, ΦPSII recovery offset, final ΦPSII and Fv/Fm were driven by data structure rather than model-specific tuning.

**Figure 3.**
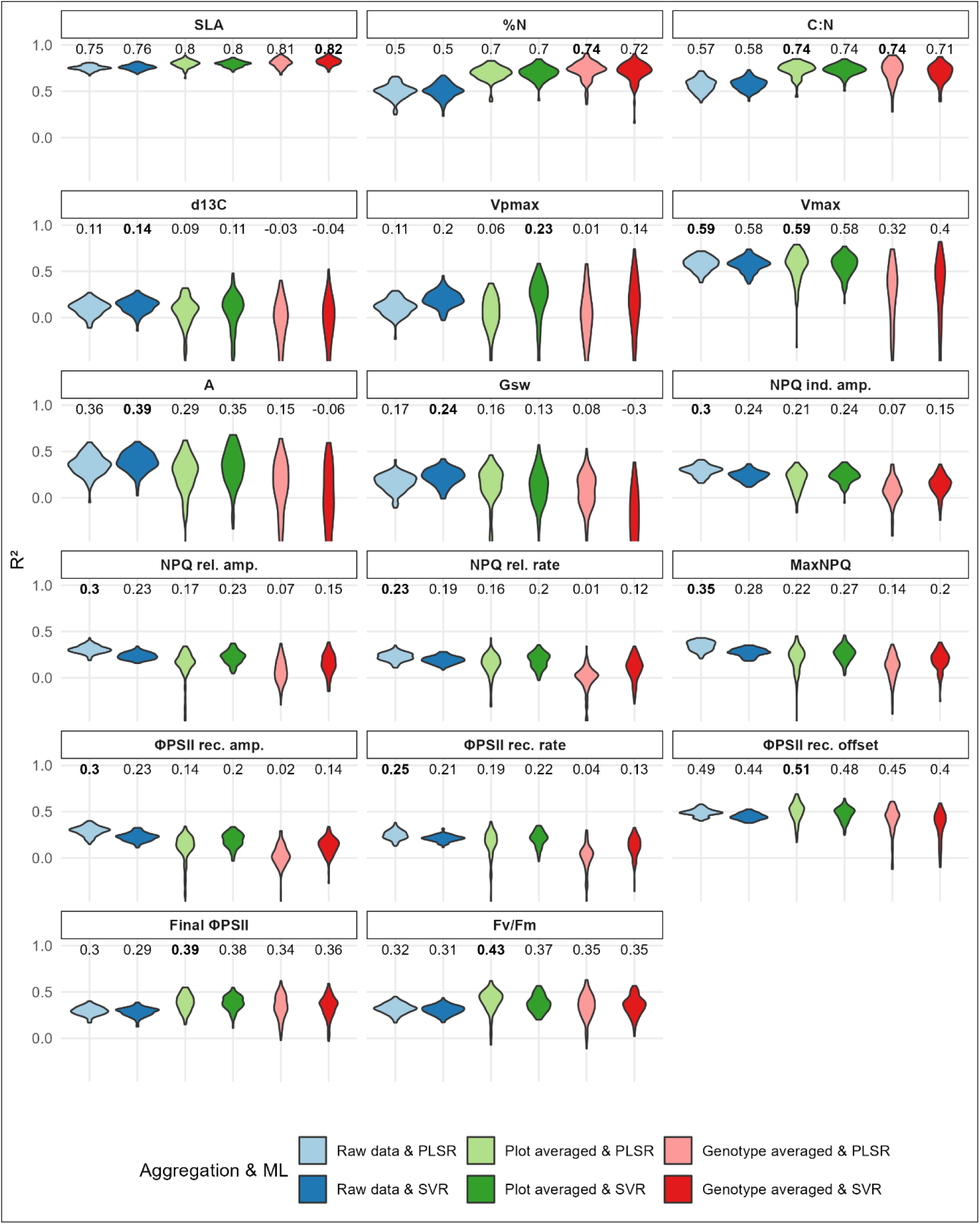
Comparison of PLSR and SVR model performance under different aggregation strategies for 17 traits measured in season 2021. Each trait was evaluated using combinations of aggregation methods and machine learning algorithms (color-coded), with model performance assessed via 20 repetitions of 5-fold CV, calibrated with single CV. Violin plots depict the distribution of R^2^ scores across 100 validation folds, with median values shown above each plot. Bolded numbers indicate the highest median R^2^ achieved among all combinations.

To further test whether SVR consistently outperform PLSR for certain traits across seasons, trait-specific aggregation strategies were employed: for SLA, %N and C:N ratio, genotype-averaged data were used; for ΦPSII recovery offset, final ΦPSII and Fv/Fm, plot-averaged data were used; for δ^13^C, even though raw data showed better performance, for season 2022 and 2023, no raw data were available, thus the comparison was made using plot-averaged data. SVR outperformed PLSR for V_pmax_, A and gsw across all three seasons while for V_max_, SVR had better performance for 2022 and 2023, and for δ^13^C and final ΦPSII was for 2021 and 2023 (Figure S7).

The previous evaluations assessed model generalizability across genotypes within single seasons. To explore whether combining data across seasons could improve predictive performance and model robustness, we tested all combinations of two and three seasons using trait-specific aggregation strategies and the best-performing models: genotype-averaged data with PLSR for SLA, %N, and C:N ratio; plot-averaged data with SVR for δ^13^C; plot-averaged data with PLSR for ΦPSII recovery offset, final ΦPSII and Fv/Fm; and raw data with SVR for the remaining traits. While several traits, including C:N ratio, NPQ relaxation amplitude, ΦPSII recovery amplitude and offset, final ΦPSII, Fv/Fm and maximum NPQ, already performed well using data from individual seasons, the combination of two or three seasons resulted in slightly improved median R^2^ for the remaining traits (Figure S8). Notably, multi-season training consistently reduced prediction uncertainty, reinforcing the advantage of integrating environmental variability to improve model robustness.

To further examine the predictive capacity of models trained using all seasons, we reconstructed full response vectors by aggregating fold-level predictions from the first CV repetition, yielding single R^2^ scores. Remarkably, models trained on combined data were able to capture differences in trait scale across seasons. For instance, SLA values from 2021 were systematically lower than in the other two seasons, and δ^13^C from 2022 exhibited a distinct distribution (Figure S9). These between-season differences were successfully captured by the ML models, demonstrating the ability of hyperspectral data to retain season-specific physiological signals and the effectiveness of multi-season models in capturing them.

Together, these results underscore the importance of tailoring data aggregation (pre-processing) strategies to the specific trait categories and compare machine learning models to optimize model performance and reliability across environmental conditions in different seasons.

### 3.4 Evaluating model generalizability to unseen genotypes and seasons

The analysis above utilized random split between calibration and validation sets, enabling a systematic evaluation of how different data aggregation strategies and seasonal variation affect the predictive performance of PLSR and SVR models. To further investigate model generalizability, we next assessed how well these models perform when predicting traits for genotypes that were not considered during calibration. This analysis was conducted using sub-sampled HSR and applied the same nested cross-validation framework within season as described above, with single CV for NoC estimation. Moreover, trait-specific ML and aggregation techniques were used as in the previous analysis. Uniquely, in this scenario, we enforced a strict separation of genotypes between the calibration and validation sets in both the inner and outer cross-validation loops, ensuring that predictions were made exclusively on unseen genotypes. This is relevant to prevent data leakage that can artificially increase model performance.

Overall, model performance for the unseen genotype scenario was consistently lower than for the random split scenario (Figure 4). For genotype-averaged structural and biochemical traits, PLSR models demonstrated strong generalizability, achieving similar R^2^ values in both the random split and unseen genotype scenarios for all three seasons. This indicates that these traits exhibit more stable relationships with HSR across genotypes. Interestingly, plot-averaged ΦPSII recovery offset, final ΦPSII and Fv/Fm showed less significant difference between random splits and predicting unseen genotypes. In contrast, gas exchange and the remaining fluorescence traits showed a marked decline in predictive performance when moving to the unseen genotype scenario.

**Figure 4.**
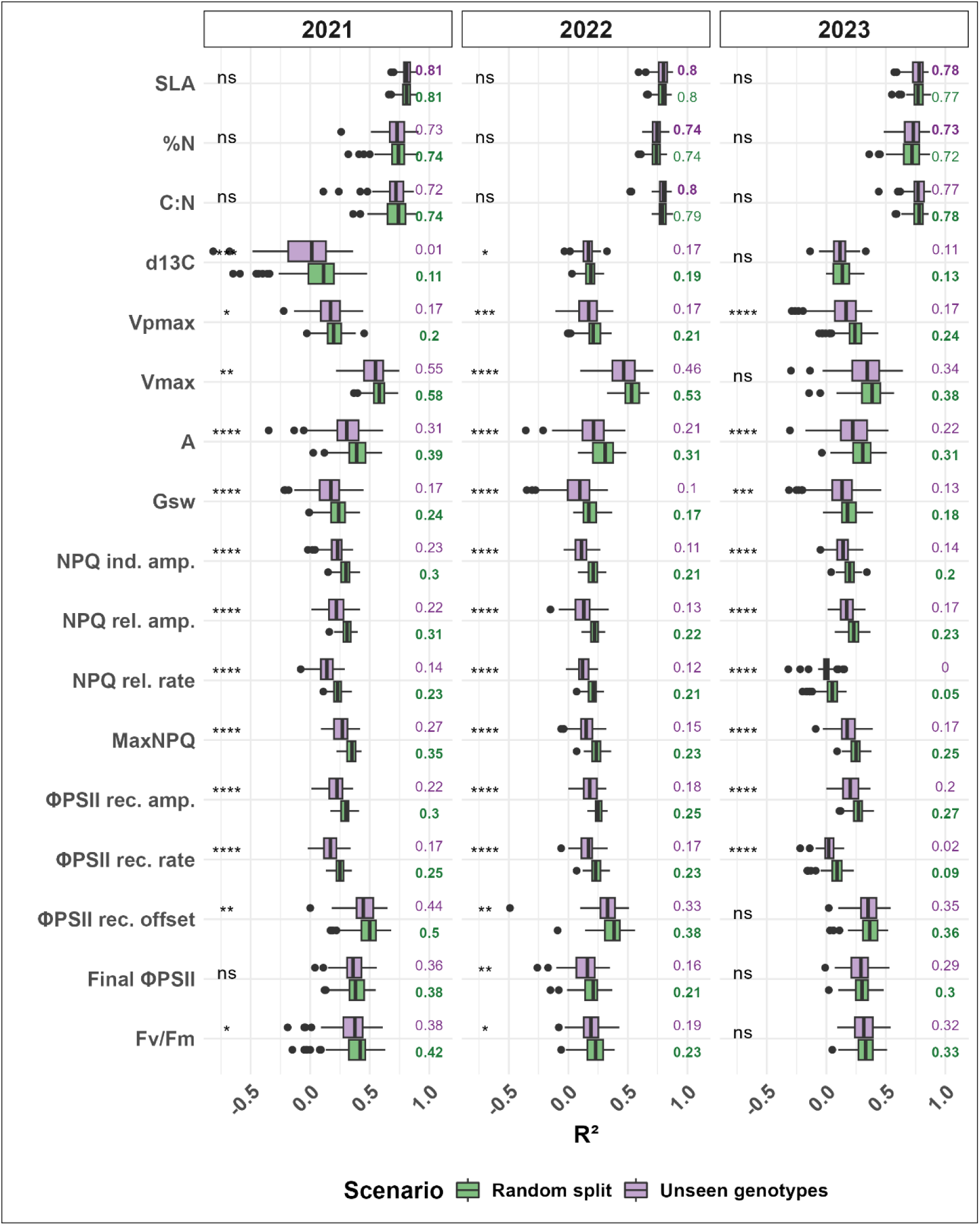
Comparison of prediction performance between random and unseen genotype scenarios using trait-specific combinations of aggregation strategies and machine learning algorithms. For 17 traits measured across three seasons, boxplots show *coefficient of determination (*R^2^) values across 20 repetitions of 5-fold CV, using the optimal combination of aggregation method and machine learning algorithm for each trait. Two prediction scenarios were evaluated: random data points splitting calibration and validation(green) and validation includes only unseen genotypes (purple). Statistical significance between scenarios is indicated by asterisks: * (p ≤ 0.05), ** (p ≤ 0.01), *** (p ≤ 0.001), **** (p ≤ 0.0001), ns = not significant.

We then expanded the analysis to evaluate model transferability across seasons by training on one or more seasons and testing on a held-out season. For instance, when predicting 2021 data, models were trained on data from 2022, 2023, or their combination, calibrated using 10 times repeated 3-fold CV for NoC estimation. Given the differences in trait magnitudes between seasons, we assessed model performance using squared Pearson correlation between predicted and observed values.

Among the three tested seasons, season 2021 consistently showed the highest predictive accuracy, particularly when trained on combined data from 2022 and 2023 data (Figure 5). In contrast, 2022 emerged as the most difficult season to predict, with limited gains based on incorporating data from both 2021 and 2023. SLA and %N were the only traits for which models trained on individual seasons produced acceptable accuracy. For predictions targeting 2023, training on the combined datasets from 2021 and 2022 resulted in the best overall performance across most traits, with the exception of SLA, where models trained on individual seasons outperformed those trained on the combined dataset. For SLA, the combination of data from seasons 2021 and 2022 and combination of data from seasons 2021 and 2023 as training set, yielded poor results. This was due to the different SLA distribution in 2021 compared to the other two seasons. To address this issue, SLA from individual seasons were scaled and centered before combining to form the training set, and the validation prediction was performed using the scaled response. The predicted values were then transformed back to the original scale. As a result, the accuracy of SLA predictions for seasons 2022 trained on combined data from seasons 2021 and 2022 increased from 0.2 to 0.60 for the squared Pearson correlation after pre-processing the genotype-averaged response variable (Figure S10). For testing season 2023, the accuracy was improved from 0.04 to 0.60. However, for the remaining traits, the effect of scaling and centering was negligible.

**Figure 5.**
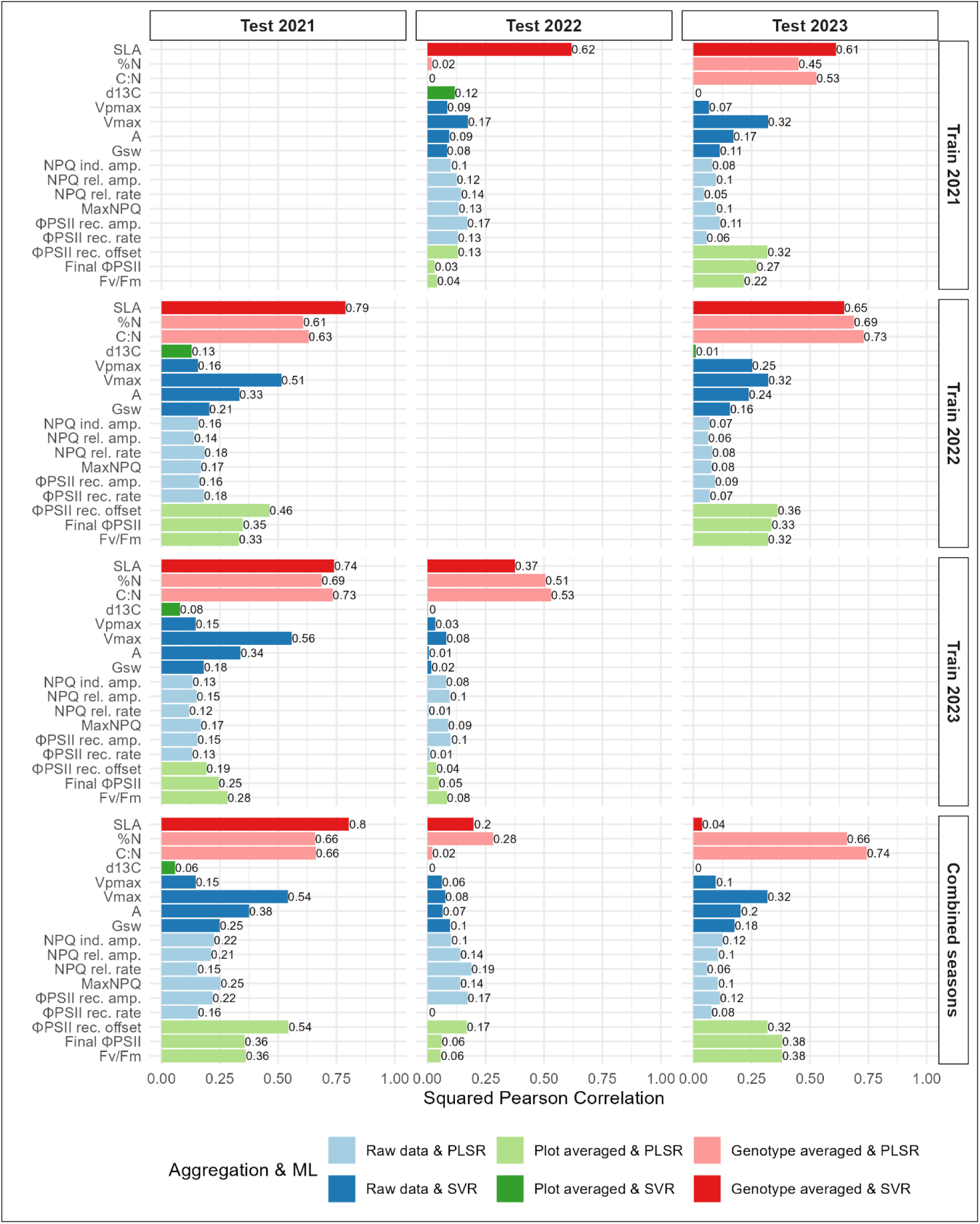
Prediction performance across unseen seasons using optimal aggregation and ML combinations. Bar plots display the squared Pearson correlation (R^2^) between predicted and measured trait values for 17 traits. Rows indicate the season(s) used to calibrate the models (2021, 2022, 2023, or combined seasons), while columns correspond to the test season used for independent validation. For each trait, models were built using the trait-specific optimal combination of aggregation strategy and machine learning algorithm (illustrated in different colors).

Another challenge included the prediction of leaf nitrogen content (%N), which exhibited poor performance when transferring models from the 2021 season to 2022—despite successful prediction in the reverse direction (from 2022 to 2021). Analysis of genotype-averaged HSR in the 1500–2400 nm range, which was used for %N prediction, revealed notable differences in spectral distributions between the two seasons (Figure S11). The genotypes exclusive to the 2022 season displayed broader spectral variation than those in 2021. Given that %N measurements in 2021 were available only for these 99 genotypes, each of which was also included in the 2022 dataset, the limited spectral variability in the 2021 training data may have limited model generalization. In support of this conclusion, Ji et al. (2024) demonstrated that greater spectral diversity in the training set enhances the predictive performance of PLSR models in new domains. Therefore, our results indicate that the reduced spectral diversity in the 2021 data contributed to the diminished transferability of %N prediction models from 2021 to 2022.

To further probe model generalizability, we evaluated the most stringent scenario: predicting traits for unseen genotypes from entirely unseen seasons. To this end, genotypes from the target season (2021) were partitioned into five folds, each fold was used for validation, while the remaining genotypes from the other two seasons formed the training set. During the inner loop for selecting NoC, genotype structure was exclusive to avoid data leakage. Across 10 repetitions for each training season (2022 and 2023), models exhibited consistently lower squared Pearson correlations than in any previous scenario (Figure S12). The performance drop from random split to unseen genotypes within the same season was smaller than the drop associated with predicting unseen genotypes for unseen seasons. Similar to the previous analysis, where the entire unseen season was predicted, here, to predict unseen genotypes from unseen season, 2022 was consistently the most challenging one.

Collectively, these results underscore the complexity of transferring hyperspectral models across genotypes and seasons. They highlight the importance of trait-specific properties, data aggregation strategies, and environmental variability in determining model robustness.

## 4. Discussion

HSR data have been used to train predictive ML models for different traits across variety of plants and crops (Kumar et al., 2001; Serbin et al., 2012; Heckmann et al., 2017; Yendrek et al., 2017; Silva-Perez et al., 2018; Cotrozzi et al., 2020; Feng et al., 2020; Furbank et al., 2021; Wang et al., 2021; Yu et al., 2022; Kaur et al., 2024). Despite the wider adoption of these data to devise ML models, several pressing issues remain poorly explored: The most important issue deals with understanding the generalizability of the ML models to data that have not been used in training. Another issue addresses the applicability of ML models with data from the field, where several data aggregation alternatives are possible. In addition, advances in ML have resulted in variety of models, each with their set of hyperparameters requiring adequate procedure for their tuning.

To address these gaps, we evaluated model generalizability across increasingly realistic prediction scenarios: (1) random data splits within season(s), (2) prediction of unseen genotypes within a season, and (3) prediction of unseen genotypes from entirely unseen seasons. Using a multi-season field dataset from a maize MAGIC population, we also explored how model choice and data aggregation influence predictive performance.

In the first part of our study, we calibrated PLSR models on the 2021 dataset, examined model sensitivity to NoC, and compared three methods for NoC estimation: repeated CV, single CV, and permutation-based selection. All approaches yielded consistent NoC estimates and similar performance on validation folds. Sub-sampling the spectral data (every fifth wavelength) provided results comparable to the full spectrum, substantially reducing computational time.

Several traits demonstrated consistently poor predictability in this calibration step, including %C, δ^15^N, iWUE, stomatal limitation, NPQ induction rate and slope, NPQ relaxation offset, and final NPQ. Testing nonlinear models (XGBoost, neural networks, SVR with radial kernel) yielded no improvement over PLSR, except for SVR with a linear kernel, which performed comparably. Consequently, for downstream analysis, we focused on comparing PLSR and linear SVR using sub-sampled spectra. Given that this analysis used the entire 2021 dataset for NoC estimation, it served as a calibration step rather than a validation of generalizability.

To properly evaluate generalizability, we employed a nested CV design, consisting of 20 outer loops of 5-fold CV. For each iteration, the inner calibration loop estimated NoC using either repeated CV or single CV, while the outer validation loop assessed model generalizability. To evaluate model generalizability using random data splits, both plot-level and genotype-level aggregation were tested across the 2021, 2022, and 2023 seasons. Weakly predictable traits identified during the calibration step remained of poor predictability regardless of pre-processing or ML approaches and were therefore excluded from further analyses.

Among structural and biochemical traits, SLA, ‰N, and C:N ratio showed strong generalizability across all scenarios for both PLSR and SVR. These traits benefited most from genotype-level averaging, which consistently improved prediction accuracy. In contrast, photosynthetic traits were best predicted using raw data combined with linear SVR. Among these, V_max_, representing the asymptotic maximum of the *A-Ci* curve under CO_2_-saturated Rubisco, demonstrated the highest generalizability. In comparison, net assimilation rate (*A*), measured at a fixed internal CO_2_ concentration of 400 ppm, may be more affected by transient environmental variability, resulting in lower predictive accuracy. The estimation of V_pmax_, estimated from the slope of the *A-Ci* response at low internal CO_2_ concentrations, showed weaker predictability, likely due to its reliance on more complex C_4_ photosynthetic mechanisms, including C_4_ acid transport, CO_2_ leakiness, and energy partitioning, which are difficult to capture through HSR alone, as previously reported (Yendrek et al., 2017).

This study is also the first to comprehensively predict a set of fluorescence-based traits using ML applied on HSR. Among these, ΦPSII recovery offset is the best predicted and reflects the quantum efficiency of PSII at the end of light exposure. Traits such as Fv/Fm (maximum PSII efficiency), final ΦPSII, and maximum NPQ showed moderate performance. Amplitude-based metrics (e.g., NPQ induction/relaxation amplitude, ΦPSII recovery amplitude) outperformed rate-based metrics, likely because amplitudes capture longer-term physiological changes detectable in reflectance, whereas rates reflect rapid processes upon light on/off switch with weak structural signatures (Ferguson et al., 2025). In general, NPQ relaxation and ΦPSII recovery traits were more predictable than NPQ induction traits, which occur rapidly and are driven by transient electrochemical gradients. In contrast, NPQ relaxation and ΦPSII recovery involve pigment transitions and structural dynamics that HSR is better suited to detect. Certain fluorescence traits also required higher NoC (∼30), suggesting these traits involve multiple pigments, biochemical reactions and dynamic regulation mechanisms, resulting in complex and distributed spectral signals.

As expected, model generalizability declined across the three scenarios, with the greatest challenge posed by predicting unseen genotypes from an entirely unseen season, the most practically relevant setting. This marked drop in performance compared to random splits underscores the influence of environmental variation on the predictive power of HSR data.

Our results demonstrate that there was no universally optimal ML model or data aggregation strategy. Instead, trait-specific approaches were necessary to reach optimal performance. HSR is well-suited for predicting integrative traits, like SLA and %N, while physiological traits, such as: ΦPSII recovery and maximum PSII efficiency, exhibited more moderate performance. Short-term, dynamic traits remain difficult to predict using HSR data.

Our work paves the way for scrutinizing the predictive power of HSR data for different classes of traits by usage of well-established CV techniques and ML models. In addition, it also points out that data transformations can have often drastic effect on model performance, particularly for prediction scenarios relevant in practice, e.g., breeding. Finally, our work reveals the persistent challenge of predicting dynamic physiological traits, motivating further research into novel feature engineering and temporal modeling approaches.

## Data availability statement

All code and raw data to ensure reproducibility of the results can be accessed at: <u>https://github.com/Rudan-X/HyperspectralML</u>

## Ethics Approval Statement

Not applicable.

## Patient Consent Statement

Not applicable.

## Permission to Reproduce Material

No third-party material was reproduced

### Abbreviations

HSR: hyperspectral reflectance
partial least squares regression: PLSR
number of component: NoC
support vector regression: SVR
*A-Ci* curve: photosynthetic rate versus internal CO_2_ concentration curve
phosphoenolpyruvate carboxylase: PEPC
maximum rate of carboxylation by PEPC: V_pmax_
asymptote of the *A-Ci* curve: V_max_
photosystem II: PSII
quantum efficiency of PSII: Fv/Fm
non-photochemical quenching: NPQ
PSII operating efficiency: ΦPSII
relative C content: %C
relative N content: %N
stable carbon isotope ratio: δ^13^C
stable nitrogen isotope ratio: δ^15^N
intrinsic water use efficiency: iWUE
stomatal conductance: gsw
coefficient of determination: R2
cross-validation: CV
coefficients of variation: CVs

---

**Figure S1.**
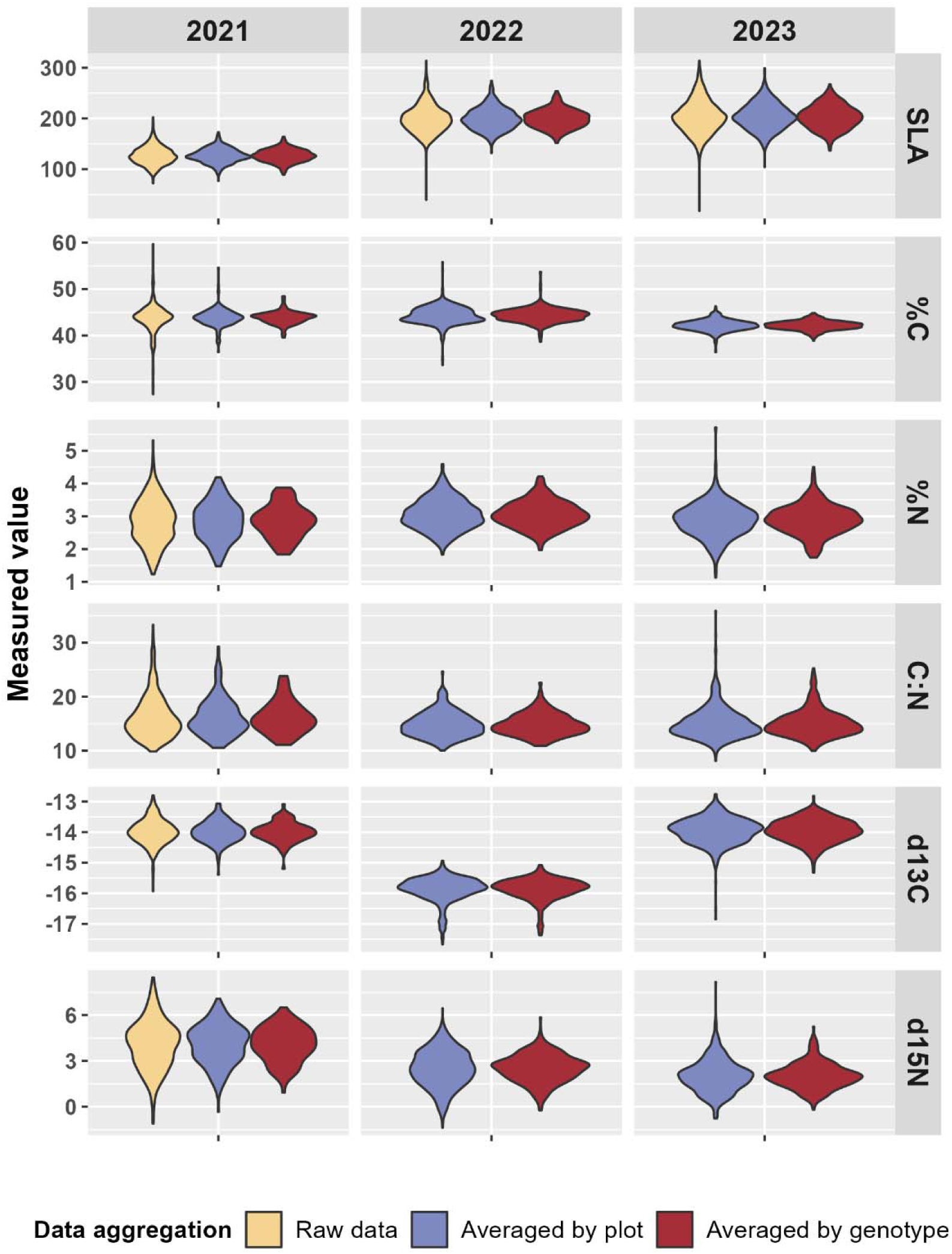
Distribution patterns of structural and biochemical traits across three consecutive growing seasons (2021–2023). Violin plots illustrate the variability and distribution of six key structural and biochemical traits: specific leaf area (SLA), percentage nitrogen (%N), percentage carbon (%C), carbon-to-nitrogen ratio (C/N), nitrogen isotope ratio (δ15N), and carbon isotope ratio (δ13C). Data are presented at three aggregation levels—raw data (yellow), averaged by plot (blue), and averaged by genotype (red)—to highlight differences in trait variability and aggregation effects over seasons.

**Figure S2.**
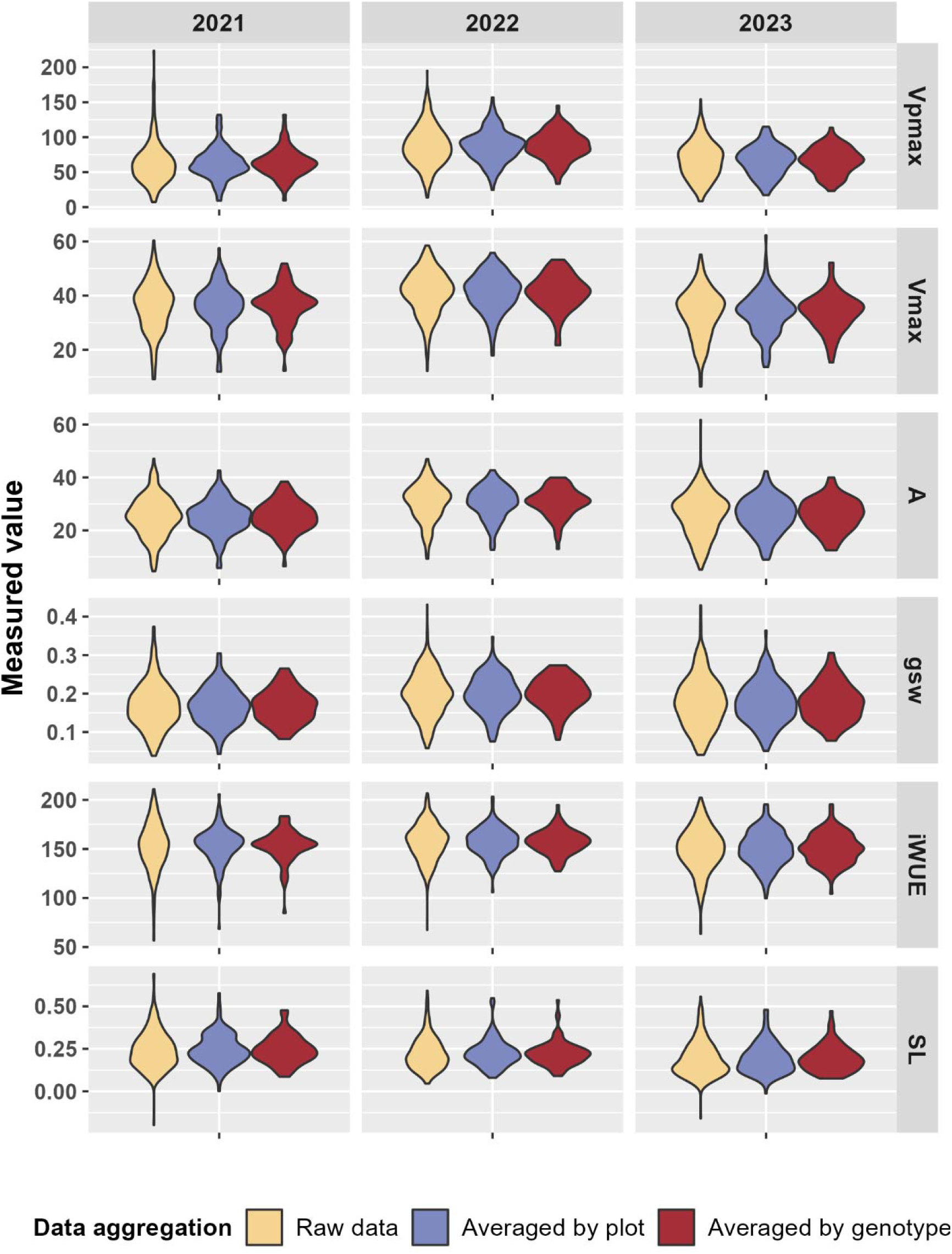
Distribution patterns of photosynthetic traits over three consecutive growing seasons (2021–2023). Violin plots illustrate variability and distributions of seven photosynthetic traits derived from gas exchange measurements: maximum carboxylation rate (V_max_), maximum electron transport rate (V_pmax_), intrinsic water-use efficiency (iWUE), stomatal conductance (Gsw), photosynthesis rate at saturating light (A), and stomatal limitation (SL) at saturating light and ambient CO^2^.

**Figure S3.**
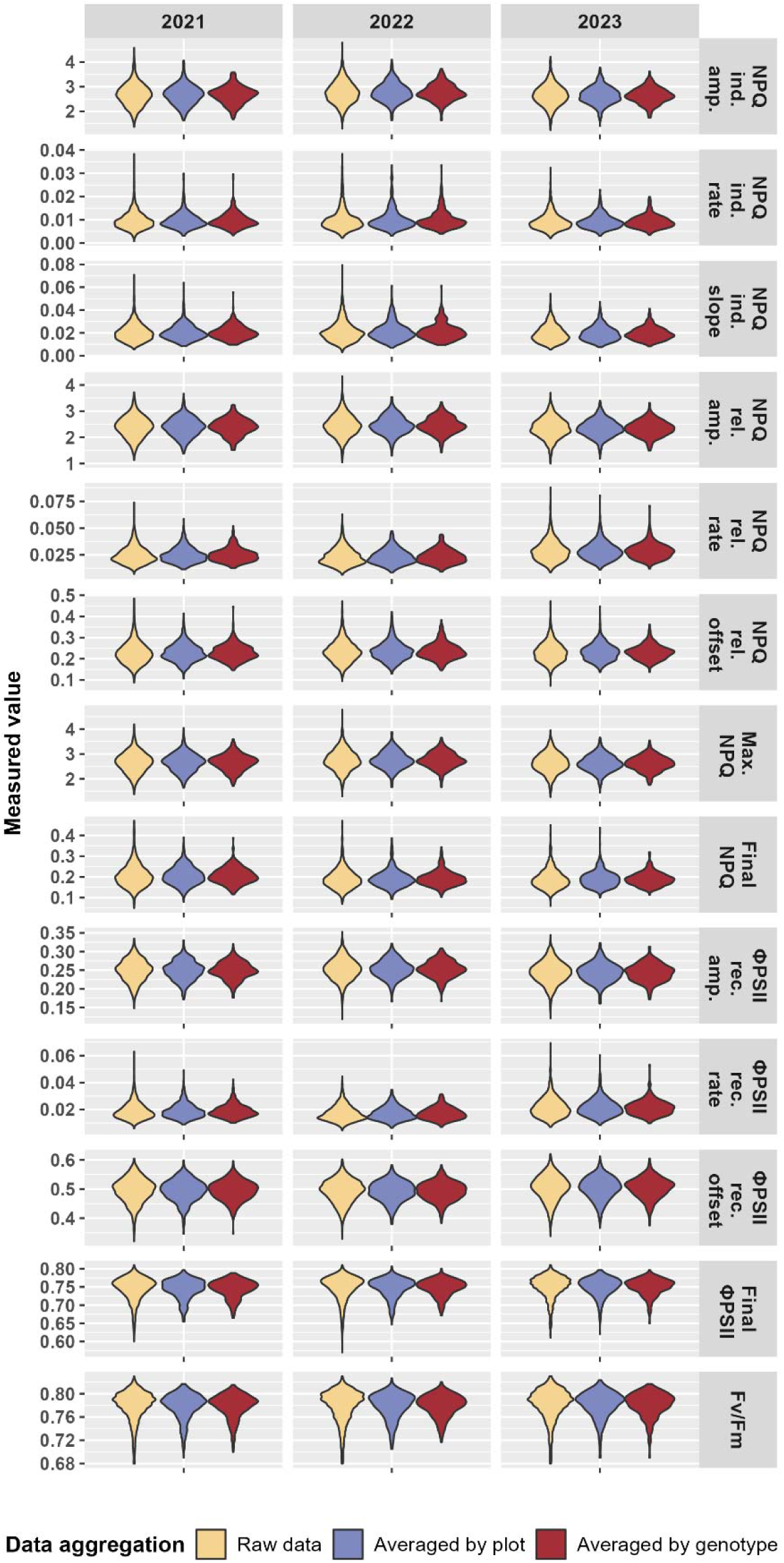
Distribution patterns of chlorophyll fluorescence traits over three consecutive growing seasons (2021–2023). Violin plots illustrate variability and distributions of thirteen chlorophyll fluorescence traits associated with photosystem II performance, including quantum efficiency (Fv/Fm), photochemical quenching efficiency (ΦPSII) and non-photochemical quenching (NPQ).

**Figure S4.**
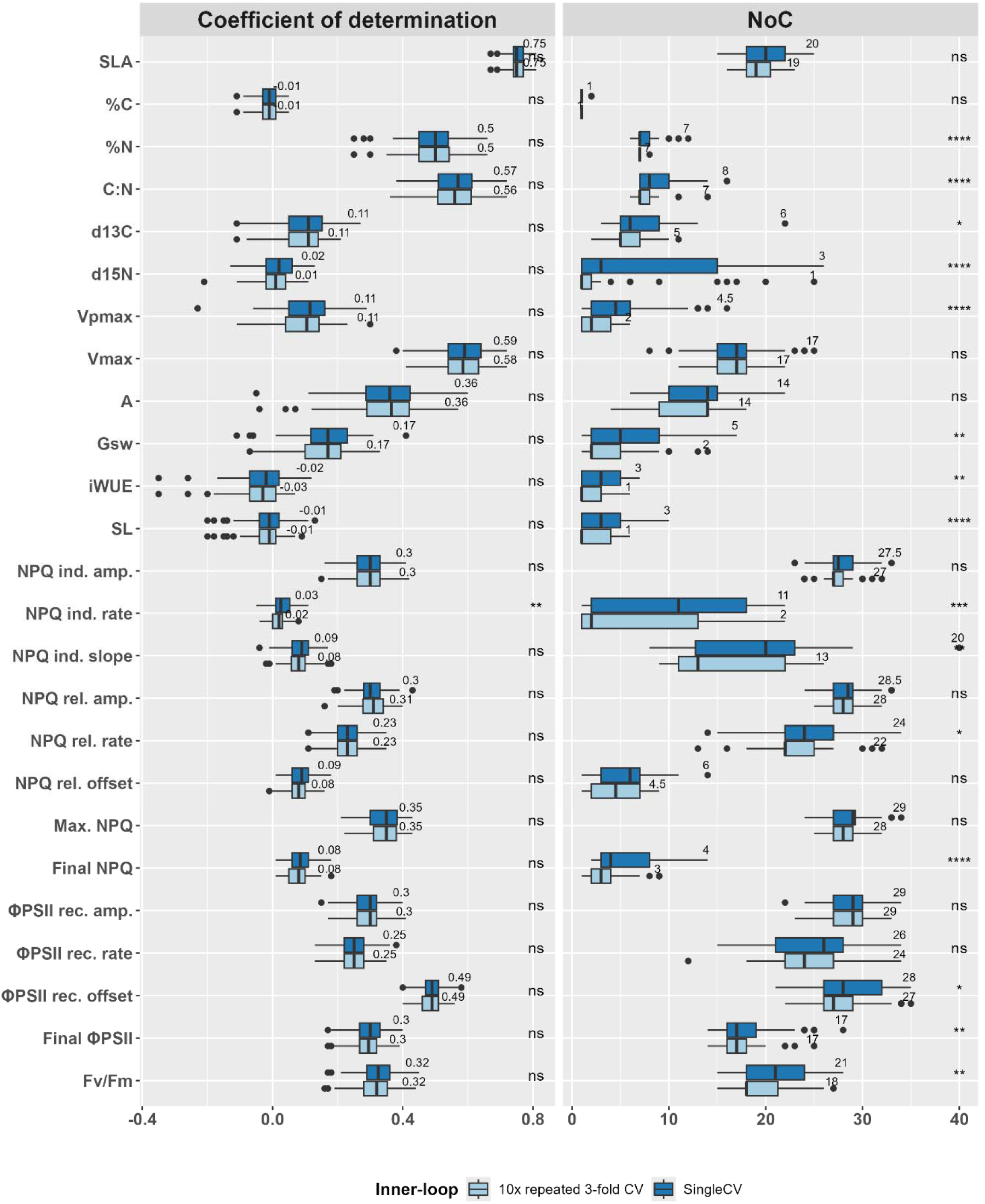
Comparison between inner-loop cross-validation strategy on model performance and optimal NoC estimation. Comparison between a single CV setup (singleCV) and 10-times repeated 3-fold CV (RepeatedCV) used in the inner loop of PLSR model tuning. Left panel shows the coefficient of determination (R^2^) for 27 traits, while the right panel shows the selected number of components (NoC) under each strategy. Boxes summarize results from 20 outer-loop repetitions of 5-fold CV. Statistical significance between the two CV strategies is indicated on the right using asterisks: * (p ≤ 0.05), ** (p ≤ 0.01), *** (p ≤ 0.001), **** (p ≤ 0.0001), ns = not significant.

**Figure S5.**
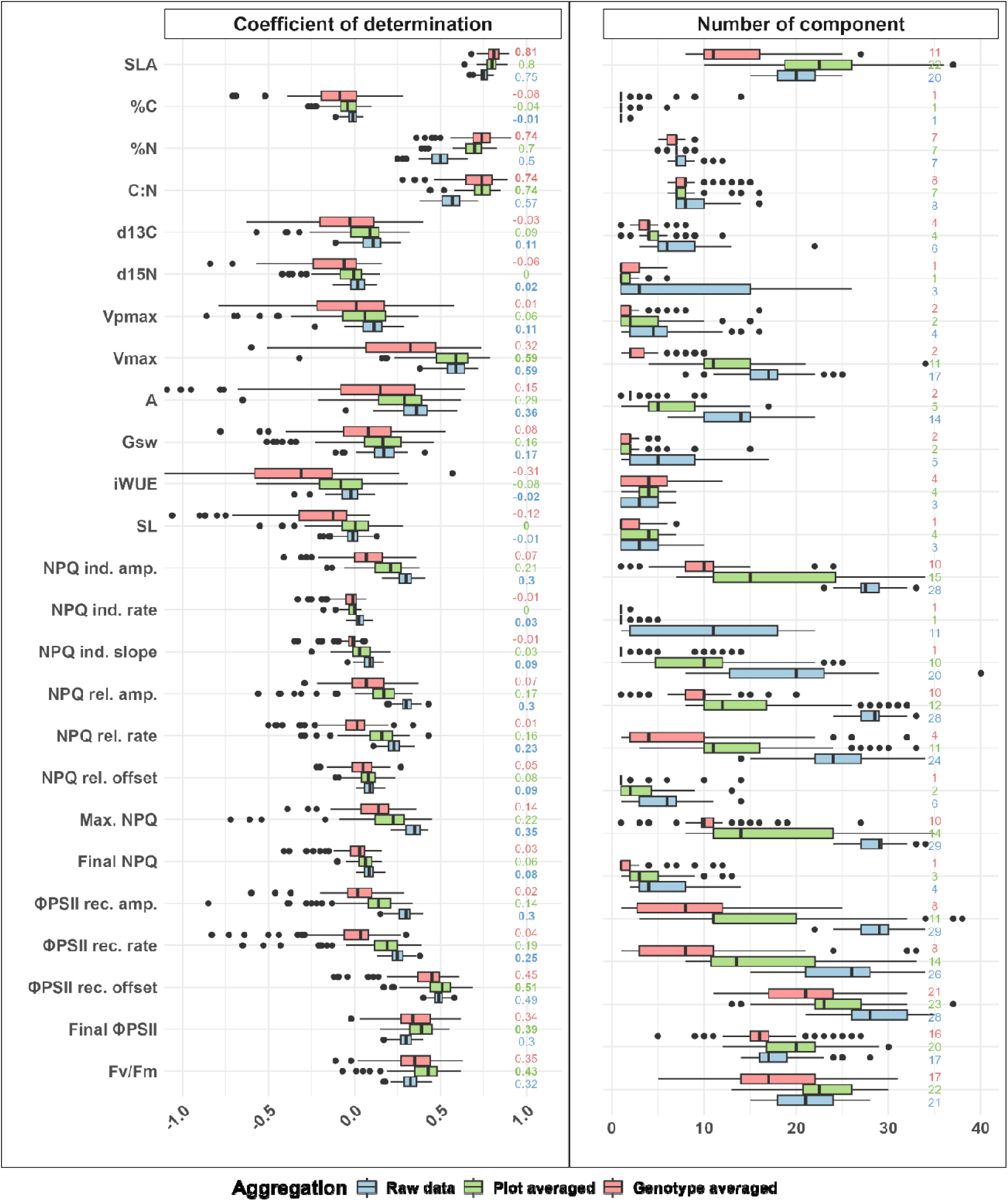
Effect of aggregation methods on model performance and selected number of components (NoC) for traits measured in the 2021 season using PLSR. The left panel shows boxplot of the coefficient of determination (R^2^) across 20 repetitions of 5-fold CV, with model tuning based on a single inner CV. The right panel displays the corresponding selected NoC in each iteration. Color indicate the three aggregation strategies: raw data (blue), plot-level averaging (green), and genotype-level averaging (red).

**Figure S6.**
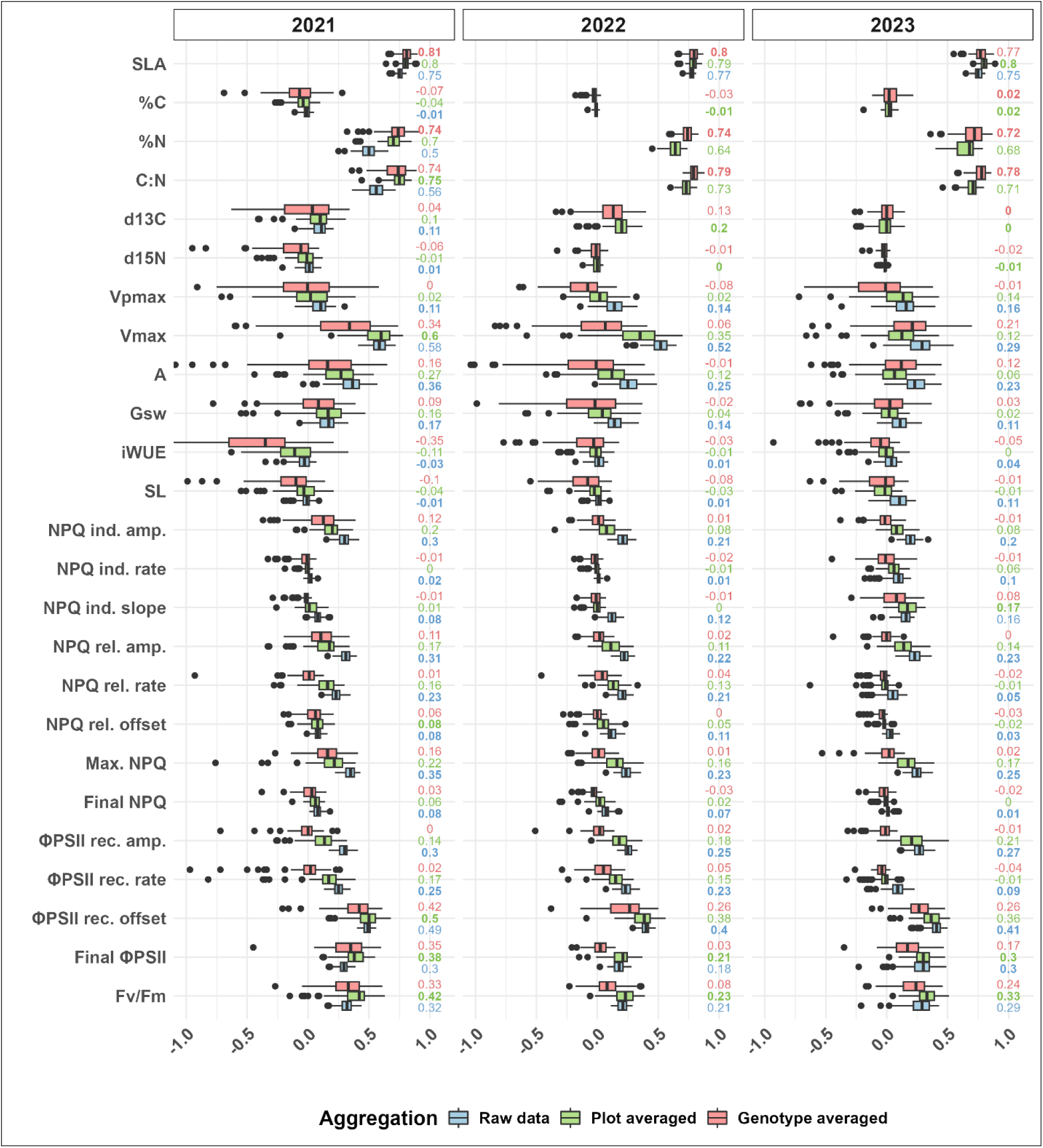
Effect of aggregation methods on model performance and selected number of components (NoC) for traits measured across three seasons using PLSR. The three panels show boxplots of the coefficient of determination (R2) for each season across 20 repetitions of 5-fold CV, with model tuning based on a single inner CV. The right panel displays the corresponding selected NoC in each iteration. Colors indicate the three aggregation strategies: raw data (blue), plot-level averaging (green), and genotype-level averaging (red).

**Figure S7.**
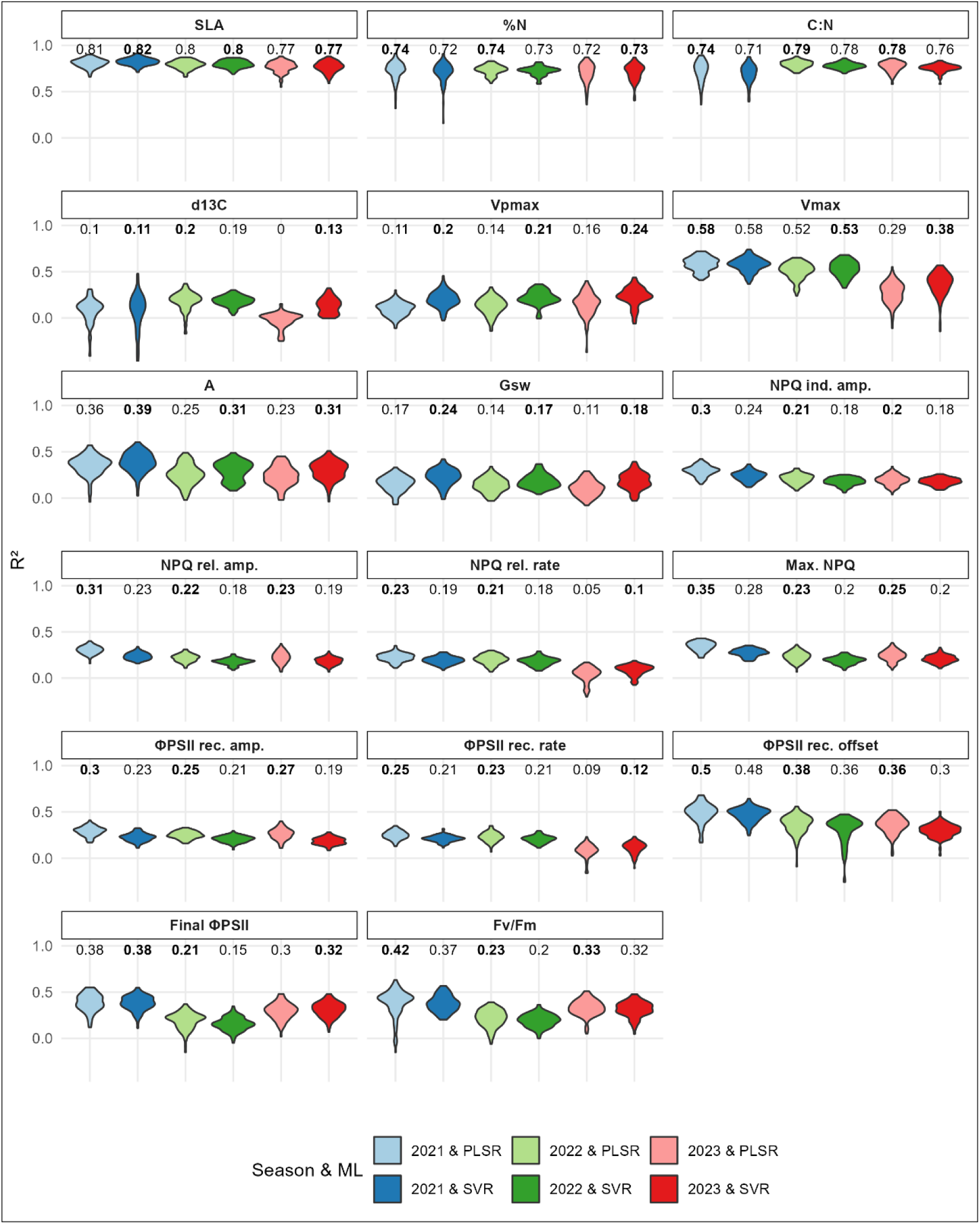
Comparison of PLSR and SVR model performance for 17 traits measured across three seasons, using optimal trait-specific aggregation method. Each trait was predicted using PLSR or SVR (light or dark colored), with model performance assessed via 20 repetitions of 5-fold CV, calibrated with single CV. Violin plots depict the distribution of R^2^ scores across 100 validation folds, with median values shown above each plot. Bolded numbers indicate the highest median R^2^ achieved among all combinations.

**Figure S8.**
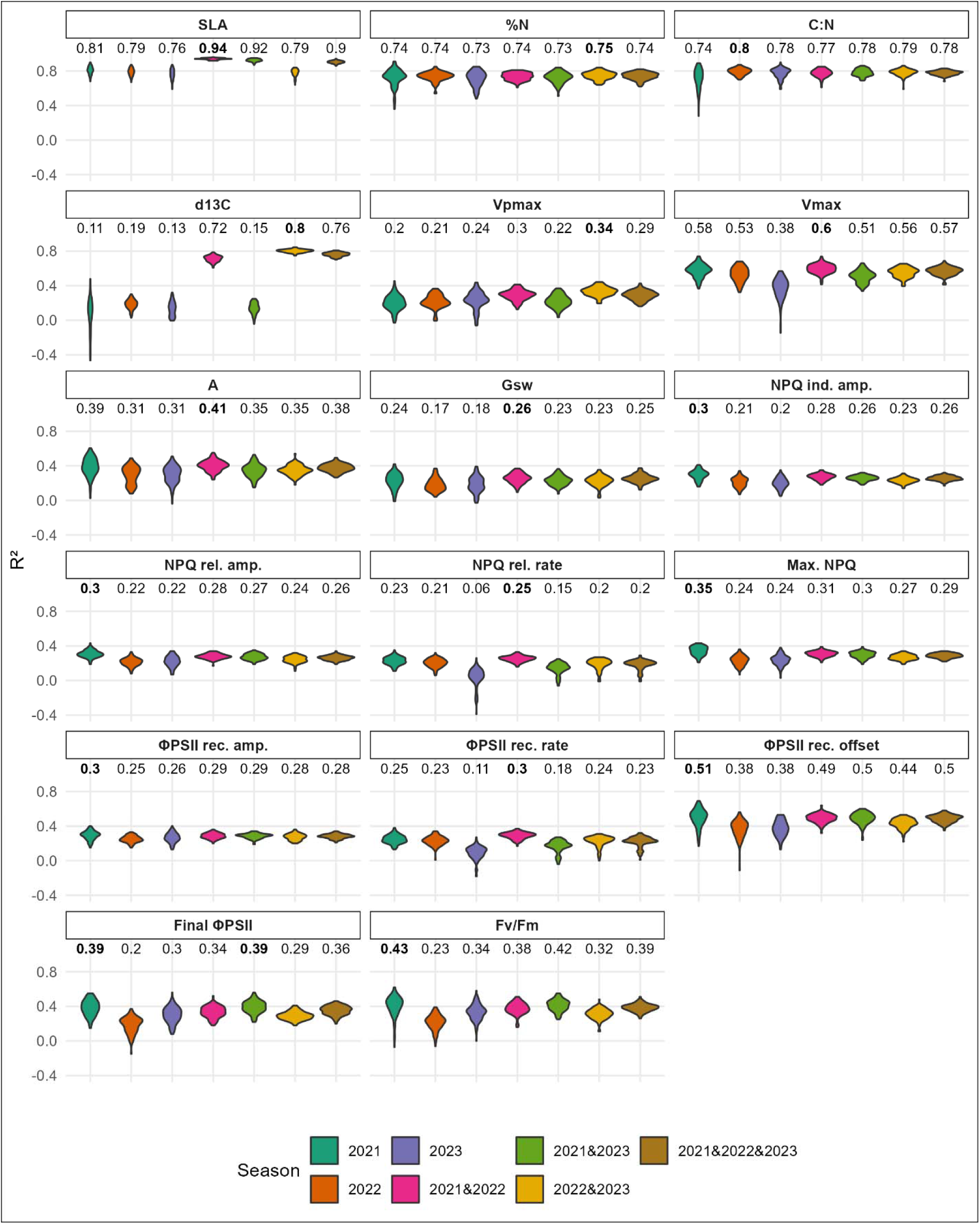
Performance of trait-specific aggregation and ML combination for individual seasons (2021, 2022, and 2023) and all combinations of these seasons. Violin plots illustrate the R^2^ distribution across 20 repetitions of 5-fold CV, with median values shown above each plot. Bolded numbers indicate the highest median R^2^ achieved among all seasons and their combinations.

**Figure S9.**
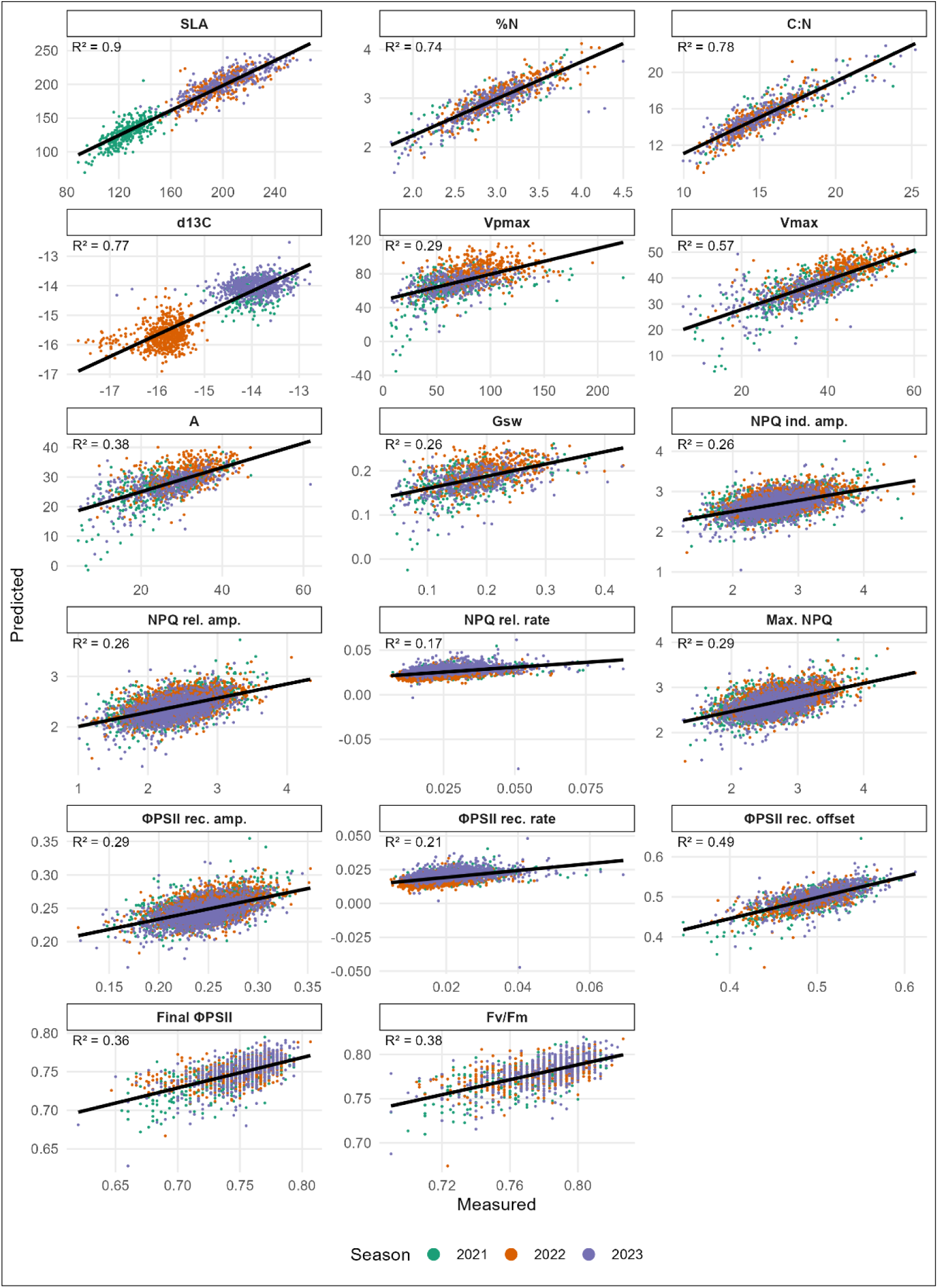
Predicted versus measured trait values from models trained on combined data from all three seasons using trait-specific aggregation and machine learning approaches. Scatterplots compare predicted values (y-axis) from the first repetition of 5-fold CV with the corresponding measured values (x-axis). Models were trained and validated using a random split strategy across combined data from the 2021, 2022, and 2023 seasons. Each subplot reports the coefficient of determination (R^2^) calculated from all test samples for the respective trait. Points are color-coded by season: green for 2021, orange for 2022, and purple for 2023.

**Figure S10.**
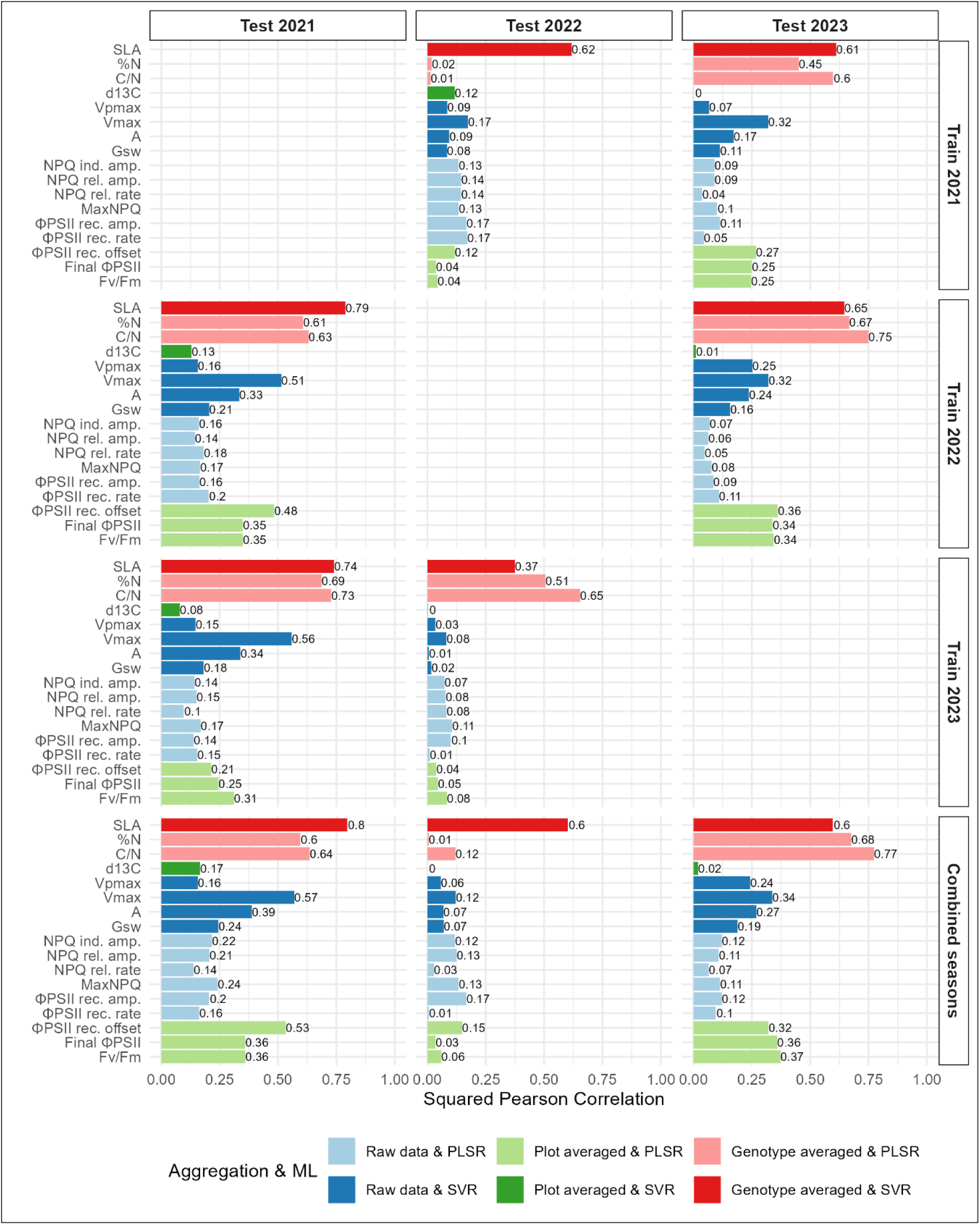
Prediction performance across unseen seasons using optimal aggregation and ML combinations, applied to scaled data. Bar plots display the squared Pearson correlation (R^2^) between predicted and measured trait values for 17 traits. Rows indicate the season(s) used to calibrate the models (2021, 2022, 2023, or combined seasons), while columns correspond to the test season used for independent validation. For each trait, models were built using the trait-specific optimal combination of aggregation strategy and machine learning algorithm (illustrated in different colors).

**Figure S11.**
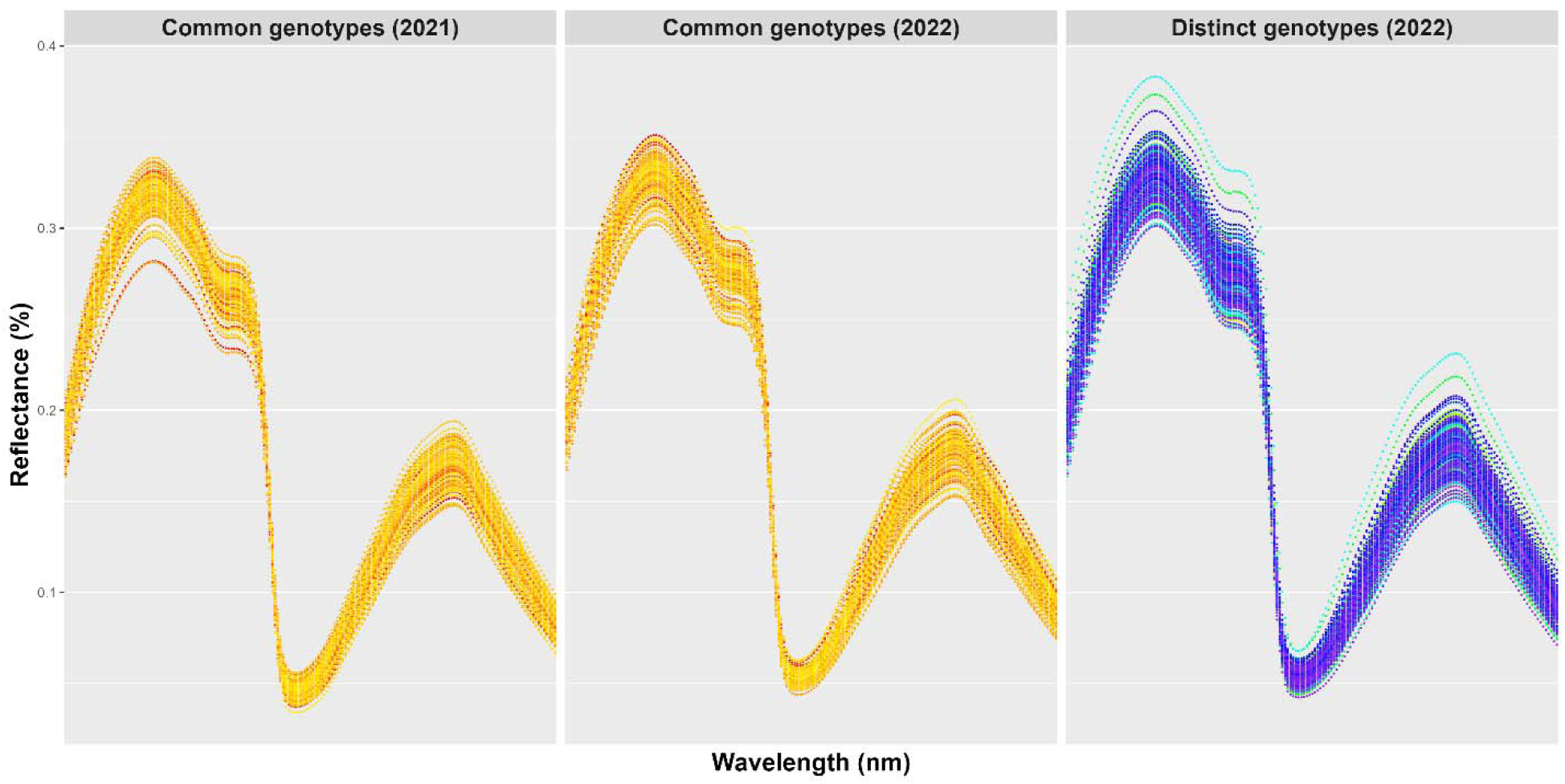
Genotype-averaged HSR profiles corresponding to %N measurements. Spectral profiles show leaf reflectance (%) across the near-infrared spectrum (1400–2400 nm). Panels differentiate 99 genotypes measured in both seasons (left and middle panels, yellow-red gradient) from 219 genotypes measured only in 2022 (right panel, blue-purple gradient). Each line corresponds to an individual leaf spectrum, illustrating the spectral variability associated with genetic and seasonal differences.

**Figure S12.**
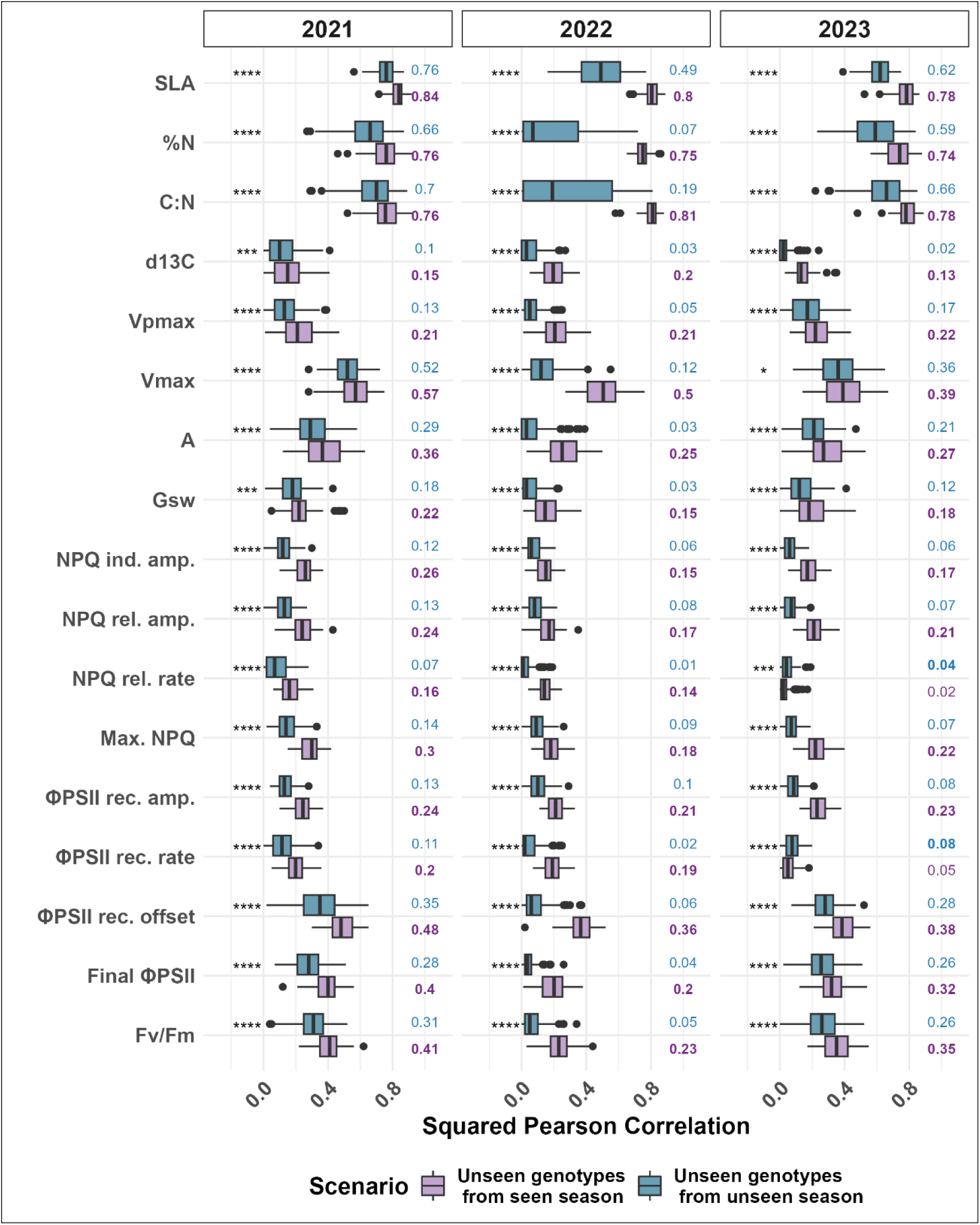
Comparison of prediction performance between unseen genotypes within the same season and unseen genotypes from unseen seasons using trait-specific aggregation and machine learning strategies. For 17 traits measured across three seasons, boxplots show square Pearson correlation values across 20 repetitions of 5-fold cross-validation, using the optimal combination of aggregation method and machine learning algorithm for each trait. Two prediction scenarios were evaluated: prediction of unseen genotypes within the same season (blue), and prediction of unseen genotypes from seasons not included in model calibration (purple). Statistical significance between scenarios was assessed using paired comparisons; significance levels are denoted as: * (p ≤ 0.05), *** (p ≤ 0.001), **** (p ≤ 0.0001).

**Table S1.**
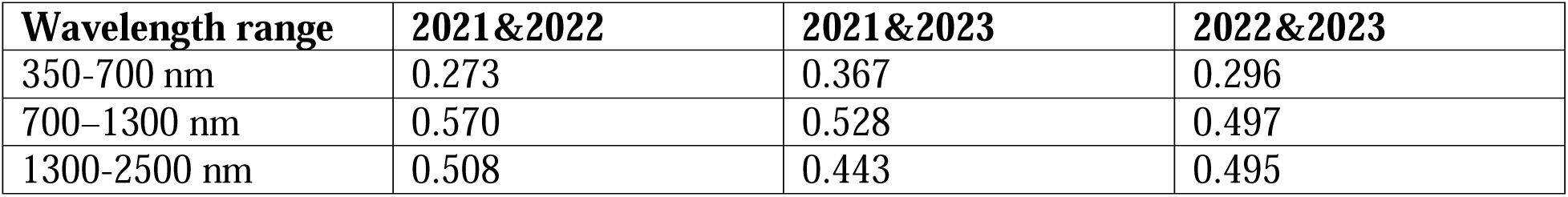
Mean correlation of HSR across different wavelengths between three seasons.

| Wavelength range | 2021&2022 | 2021&2023 | 2022&2023 |
| --- | --- | --- | --- |
| 350-700 nm | 0.273 | 0.367 | 0.296 |
| 700–1300 nm | 0.570 | 0.528 | 0.497 |
| 1300-2500 nm | 0.508 | 0.443 | 0.495 |

**Table S2.** Number of raw data samples collected over three consecutive seasons. The values in parentheses represent the number of samples aggregated by averaging per plot and per genotype.

| Trait | 2021 | 2022 | 2023 |
| --- | --- | --- | --- |
| SLA | 1845 (623,316) | 1895 (633,318) | 1669 (556,310) |
| C, N and isotopes | 600 (201,102) | 1069 (624,319) | 936 (618,320) |
| Chlorophyll fluorescence | 1810 (626,316) | 1678 (557,312) | 3312 (586,318) |
| Gas exchange | 382 (141,78) | 441 (161,88) | 373 (164,91) |

**Table S3.** Pearson correlation of genotype-averaged traits across three seasons.

| **Trait** | **2021&2022** | **2021&2023** | **2022&2023** | **Mean correlation** |
| --- | --- | --- | --- | --- |
| SLA | 0.473 | 0.45 | 0.361 | 0.428 |
| %C | -0.112 | 0.196 | 0.037 | 0.04 |
| %N | 0.551 | 0.441 | 0.565 | 0.519 |
| C:N | 0.568 | 0.443 | 0.571 | 0.527 |
| d13C | 0.543 | 0.555 | 0.465 | 0.521 |
| d15N | 0.245 | 0.092 | 0.283 | 0.207 |
| Vpmax | 0.212 | 0.392 | 0.35 | 0.318 |
| Vmax | 0.558 | 0.474 | 0.374 | 0.468 |
| Photosynthetic rate (A) | 0.463 | 0.416 | 0.393 | 0.424 |
| Stomatal conductance (Gsw) | 0.488 | 0.29 | 0.349 | 0.375 |
| iWUE (A/gsw) | 0.264 | 0.219 | 0.245 | 0.243 |
| Stomatal limitation | 0.566 | 0.329 | 0.556 | 0.484 |
| NPQ induction amplitude | 0.537 | 0.59 | 0.511 | 0.546 |
| NPQ induction rate | 0.423 | 0.568 | 0.344 | 0.445 |
| NPQ induction slope | 0.526 | 0.549 | 0.419 | 0.498 |
| NPQ relaxation amplitude | 0.604 | 0.635 | 0.55 | 0.596 |
| NPQ relaxation rate | 0.582 | 0.551 | 0.454 | 0.529 |
| NPQ relaxation offset | 0.32 | 0.301 | 0.356 | 0.326 |
| Maximum NPQ | 0.585 | 0.607 | 0.535 | 0.576 |
| Final NPQ | 0.306 | 0.37 | 0.28 | 0.319 |
| ΦPSII recovery amplitude | 0.634 | 0.658 | 0.619 | 0.637 |
| ΦPSII recovery rate | 0.599 | 0.552 | 0.452 | 0.534 |
| ΦPSII recovery offset | 0.6 | 0.616 | 0.654 | 0.623 |
| Final ΦPSII | 0.429 | 0.482 | 0.441 | 0.451 |
| Fv/Fm | 0.475 | 0.521 | 0.487 | 0.495 |

**Table S4.**
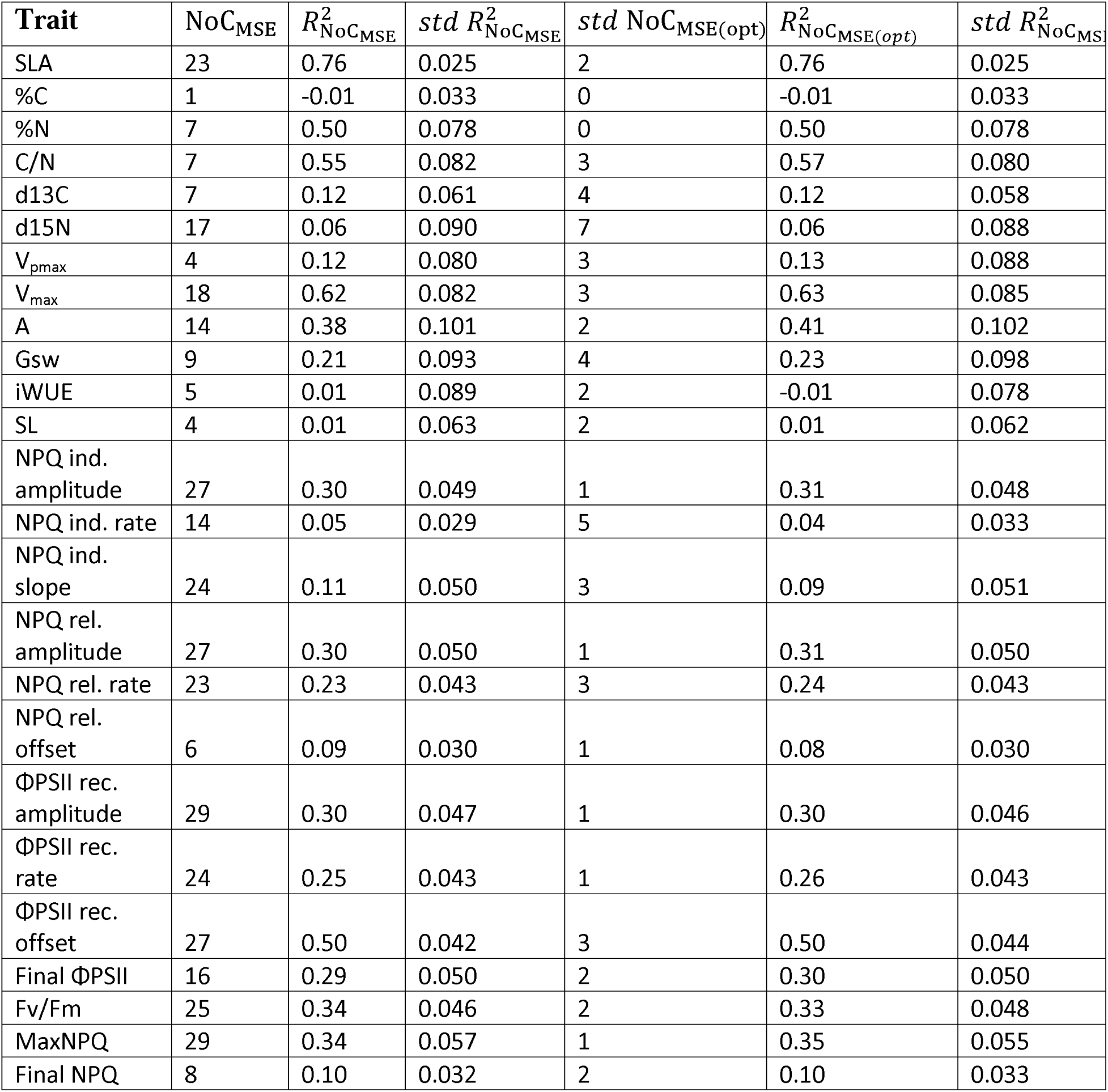
Statistics of estimated optimal NoC for each repetition of 5-fold CV using MSE-based method (. *NoC_MSE(opt)_*) and final NoC across 20 repetitions (*NoC_MSE_*) using the dataset from season 2021 as calibration data. The median R^2^ was computed across all 20 × 5 test folds using either *NoC_MSE(opt)_* (*R^2^_NOCMSE_*) or *NoC_MSE_* (*R^2^ _NOCMSE(opt)_*).

| Trait | $NoC_{MSE}$ | $R^2_{NoC_{MSE}}$ | $std R^2_{NoC_{MSE}}$ | $std NoC_{MSE(opt)}$ | $R^2_{NoC_{MSE(opt)}}$ | $std R^2_{NoC_{MSE(opt)}}$ |
| --- | --- | --- | --- | --- | --- | --- |
| SLA | 23 | 0.76 | 0.025 | 2 | 0.76 | 0.025 |
| %C | 1 | -0.01 | 0.033 | 0 | -0.01 | 0.033 |
| %N | 7 | 0.50 | 0.078 | 0 | 0.50 | 0.078 |
| C/N | 7 | 0.55 | 0.082 | 3 | 0.57 | 0.080 |
| d13C | 7 | 0.12 | 0.061 | 4 | 0.12 | 0.058 |
| d15N | 17 | 0.06 | 0.090 | 7 | 0.06 | 0.088 |
| $V_{pmax}$ | 4 | 0.12 | 0.080 | 3 | 0.13 | 0.088 |
| $V_{max}$ | 18 | 0.62 | 0.082 | 3 | 0.63 | 0.085 |
| A | 14 | 0.38 | 0.101 | 2 | 0.41 | 0.102 |
| Gsw | 9 | 0.21 | 0.093 | 4 | 0.23 | 0.098 |
| iWUE | 5 | 0.01 | 0.089 | 2 | -0.01 | 0.078 |
| SL | 4 | 0.01 | 0.063 | 2 | 0.01 | 0.062 |
| NPQ ind. amplitude | 27 | 0.30 | 0.049 | 1 | 0.31 | 0.048 |
| NPQ ind. rate | 14 | 0.05 | 0.029 | 5 | 0.04 | 0.033 |
| NPQ ind. slope | 24 | 0.11 | 0.050 | 3 | 0.09 | 0.051 |
| NPQ rel. amplitude | 27 | 0.30 | 0.050 | 1 | 0.31 | 0.050 |
| NPQ rel. rate | 23 | 0.23 | 0.043 | 3 | 0.24 | 0.043 |
| NPQ rel. offset | 6 | 0.09 | 0.030 | 1 | 0.08 | 0.030 |
| $\Phi PSII$ rec. amplitude | 29 | 0.30 | 0.047 | 1 | 0.30 | 0.046 |
| $\Phi PSII$ rec. rate | 24 | 0.25 | 0.043 | 1 | 0.26 | 0.043 |
| $\Phi PSII$ rec. offset | 27 | 0.50 | 0.042 | 3 | 0.50 | 0.044 |
| Final $\Phi PSII$ | 16 | 0.29 | 0.050 | 2 | 0.30 | 0.050 |
| Fv/Fm | 25 | 0.34 | 0.046 | 2 | 0.33 | 0.048 |
| MaxNPQ | 29 | 0.34 | 0.057 | 1 | 0.35 | 0.055 |
| Final NPQ | 8 | 0.10 | 0.032 | 2 | 0.10 | 0.033 |

**Table S5.** Statistics of estimated optimal NoC for 20 repetitions of 5-fold CV using PRESS-based( *NoC_PRESS_*) using the dataset from season 2021 as calibration data. The median R^2^ was computed across all 20 × 5 test folds using *NoC_PRESS_* (*R*^2^*_NOCPRESS_*).

| **Trait** | $NoC_{PRESS}$ | $R^2_{NoC_{PRESS}}$ | $std R^2_{NoC_{PRESS}}$ |
| --- | --- | --- | --- |
| SLA | 19 | 0.76 | 0.027 |
| %C | 1 | -0.01 | 0.034 |
| %N | 7 | 0.51 | 0.07 |
| C/N | 8 | 0.57 | 0.085 |
| d13C | 6 | 0.11 | 0.062 |
| d15N | 17 | 0.09 | 0.082 |
| $V_{pmax}$ | 4 | 0.11 | 0.087 |
| $V_{max}$ | 18 | 0.62 | 0.076 |
| A | 14 | 0.40 | 0.094 |
| Gsw | 9 | 0.22 | 0.100 |
| iWUE | 3 | 0.00 | 0.066 |
| SL | 3 | 0.00 | 0.050 |
| NPQ ind. amplitude | 27 | 0.29 | 0.051 |
| NPQ ind. rate | 13 | 0.04 | 0.028 |
| NPQ ind. slope | 20 | 0.10 | 0.050 |
| NPQ rel. amplitude | 29 | 0.30 | 0.052 |
| NPQ rel. rate | 23 | 0.23 | 0.045 |
| NPQ rel. offset | 5 | 0.09 | 0.031 |
| $\Phi PSII$ rec. amplitude | 29 | 0.31 | 0.051 |
| $\Phi PSII$ rec. rate | 24 | 0.26 | 0.047 |
| $\Phi PSII$ rec. offset | 27 | 0.50 | 0.040 |
| Final $\Phi PSII$ | 19 | 0.32 | 0.050 |
| Fv/Fm | 21 | 0.32 | 0.052 |
| MaxNPQ | 28 | 0.35 | 0.051 |
| Final NPQ | 4 | 0.08 | 0.029 |

**Table S6.**
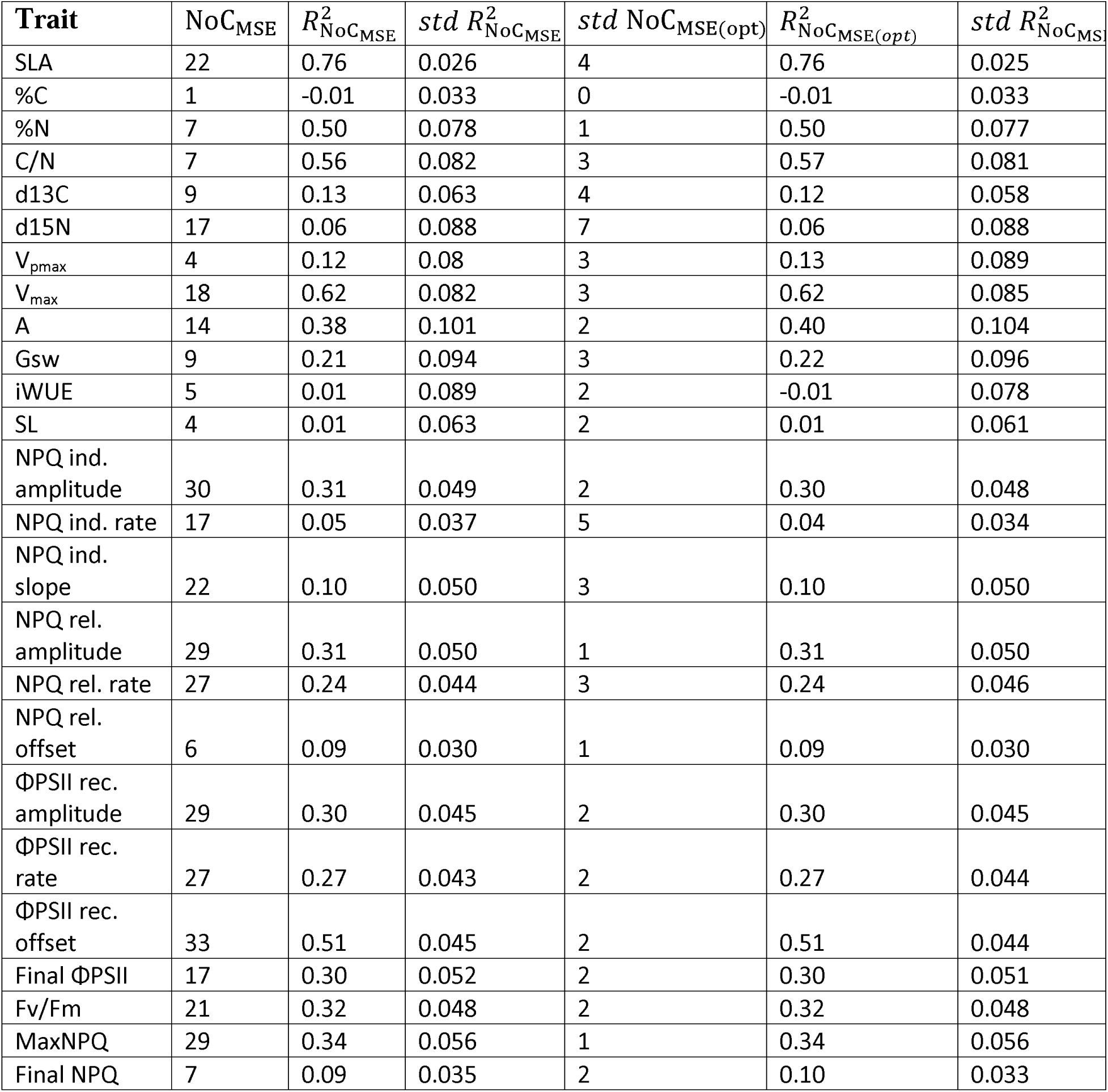
Statistics of estimated optimal NoC for each repetition of 5-fold CV (*NoC_MSE(opt)_*) and final NoC across 20 repetitions (*NoC_MSE_*) using the dataset from season 2021 as calibration data, with sub-sampled HSR data as features. The median R^2^ was computed across all 20 × 5 test folds using either *NoC_MSE(opt)_* (*R*^2^*_NOCMES_*) or *NoC_MES_* (*R^2^_NOCMSE(opt)_*).

| **Trait** | $NoC_{MSE}$ | $R^2_{NoC_{MSE}}$ | $std R^2_{NoC_{MSE}}$ | $std NoC_{MSE(opt)}$ | $R^2_{NoC_{MSE(opt)}}$ | $std R^2_{NoC_{MSE(opt)}}$ |
| --- | --- | --- | --- | --- | --- | --- |
| SLA | 22 | 0.76 | 0.026 | 4 | 0.76 | 0.025 |
| %C | 1 | -0.01 | 0.033 | 0 | -0.01 | 0.033 |
| %N | 7 | 0.50 | 0.078 | 1 | 0.50 | 0.077 |
| C/N | 7 | 0.56 | 0.082 | 3 | 0.57 | 0.081 |
| d13C | 9 | 0.13 | 0.063 | 4 | 0.12 | 0.058 |
| d15N | 17 | 0.06 | 0.088 | 7 | 0.06 | 0.088 |
| $V_{pmax}$ | 4 | 0.12 | 0.08 | 3 | 0.13 | 0.089 |
| $V_{max}$ | 18 | 0.62 | 0.082 | 3 | 0.62 | 0.085 |
| A | 14 | 0.38 | 0.101 | 2 | 0.40 | 0.104 |
| Gsw | 9 | 0.21 | 0.094 | 3 | 0.22 | 0.096 |
| iWUE | 5 | 0.01 | 0.089 | 2 | -0.01 | 0.078 |
| SL | 4 | 0.01 | 0.063 | 2 | 0.01 | 0.061 |
| NPQ ind. amplitude | 30 | 0.31 | 0.049 | 2 | 0.30 | 0.048 |
| NPQ ind. rate | 17 | 0.05 | 0.037 | 5 | 0.04 | 0.034 |
| NPQ ind. slope | 22 | 0.10 | 0.050 | 3 | 0.10 | 0.050 |
| NPQ rel. amplitude | 29 | 0.31 | 0.050 | 1 | 0.31 | 0.050 |
| NPQ rel. rate | 27 | 0.24 | 0.044 | 3 | 0.24 | 0.046 |
| NPQ rel. offset | 6 | 0.09 | 0.030 | 1 | 0.09 | 0.030 |
| $\Phi PSII$ rec. amplitude | 29 | 0.30 | 0.045 | 2 | 0.30 | 0.045 |
| $\Phi PSII$ rec. rate | 27 | 0.27 | 0.043 | 2 | 0.27 | 0.044 |
| $\Phi PSII$ rec. offset | 33 | 0.51 | 0.045 | 2 | 0.51 | 0.044 |
| Final $\Phi PSII$ | 17 | 0.30 | 0.052 | 2 | 0.30 | 0.051 |
| Fv/Fm | 21 | 0.32 | 0.048 | 2 | 0.32 | 0.048 |
| MaxNPQ | 29 | 0.34 | 0.056 | 1 | 0.34 | 0.056 |
| Final NPQ | 7 | 0.09 | 0.035 | 2 | 0.10 | 0.033 |

## References

1. Alseekh, S., Kostova, D., Bulut, M., & Fernie, A. R. (2021). Genome-wide association studies: Assessing trait characteristics in model and crop plants. Cellular and Molecular Life Sciences, 78, 5743–5754.

2. Burnett, A. C., Anderson, J., Davidson, K. J., Ely, K. S., Lamour, J., Li, Q., Morrison, B. D., Yang, D., Rogers, A., & Serbin, S. P. (2021). A best-practice guide to predicting plant traits from leaf-level hyperspectral data using partial least squares regression. Journal of Experimental Botany, 72(18), 6175–6189.

3. Chen, S., Hong, X., Harris, C. J., & Sharkey, P. M. (2004). Sparse modeling using orthogonal forward regression with press statistic and regularization. IEEE Transactions on Systems, Man, and Cybernetics, Part B (Cybernetics), 34(2), 898–911.

4. Cotrozzi, L., Peron, R., Tuinstra, M. R., Mickelbart, M. V., & Couture, J. J. (2020). Spectral phenotyping of physiological and anatomical leaf traits related with maize water status. Plant physiology, 184(3), 1363–1377.

5. Dechant, B., Cuntz, M., Vohland, M., Schulz, E., & Doktor, D. (2017). Estimation of photosynthesis traits from leaf reflectance spectra: Correlation to nitrogen content as the dominant mechanism. Remote Sensing of Environment, 196, 279–292.

6. Dell’Acqua, M., Gatti, D. M., Pea, G., Cattonaro, F., Coppens, F., Magris, G., Hlaing, A. L., Aung, H. H., Nelissen, H., Baute, J., et al. (2015). Genetic properties of the magic maize population: A new platform for high definition qtl mapping in zea mays. Genome biology, 16, 1–23.

7. Ely, K. S., Burnett, A. C., Lieberman-Cribbin, W., Serbin, S. P., & Rogers, A. (2019). Spectroscopy can predict key leaf traits associated with source–sink balance and carbon– nitrogen status. Journal of experimental botany, 70(6), 1789–1799.

8. Feng, X., Zhan, Y., Wang, Q., Yang, X., Yu, C., Wang, H., Tang, Z., Jiang, D., Peng, C., & He, Y. (2020). Hyperspectral imaging combined with machine learning as a tool to obtain high-throughput plant salt-stress phenotyping. The Plant Journal, 101(6), 1448–1461.

9. Ferguson, J. N., Caproni, L., Walter, J., Shaw, K., Arce-Cubas, L., Baines, A., Thein, M. S., Mager, S., Taylor, G., Cackett, L., et al. (2025). A deficient cp24 allele defines variation for dynamic nonphotochemical quenching and photosystem ii efficiency in maize. The Plant Cell, 37(4), koaf063.

10. Ferguson, J. N., Jithesh, T., Lawson, T., & Kromdijk, J. (2023). Excised leaves show limited and species-specific effects on photosynthetic parameters across crop functional types. Journal of Experimental Botany, 74(21), 6662–6676.

11. Ferguson, J. N., Schmuker, P., Dmitrieva, A., Quach, T., Zhang, T., Ge, Z., Nersesian, N., Sato, S. J., Clemente, T. E., & Leakey, A. D. (2024). Reducing stomatal density by expression of a synthetic epidermal patterning factor increases leaf intrinsic water use efficiency and reduces plant water use in a c4 crop. Journal of experimental botany, erae289.

12. Filzmoser, P., Liebmann, B., & Varmuza, K. (2009). Repeated double cross validation. Journal of Chemometrics: A Journal of the Chemometrics Society, 23(4), 160–171.

13. Furbank, R. T., Silva-Perez, V., Evans, J. R., Condon, A. G., Estavillo, G. M., He, W., Newman, S., Poiré, R., Hall, A., & He, Z. (2021). Wheat physiology predictor: Predicting physiological traits in wheat from hyperspectral reflectance measurements using deep learning. Plant Methods, 17, 1–15.

14. Geladi, P., & Kowalski, B. R. (1986). Partial least-squares regression: A tutorial. Analytica chimica acta, 185, 1–17.

15. Grzybowski, M., Wijewardane, N. K., Atefi, A., Ge, Y., & Schnable, J. C. (2021). Hyperspectral reflectance-based phenotyping for quantitative genetics in crops: Progress and challenges. Plant Communications, 2(4).

16. Heckmann, D., Schlüter, U., & Weber, A. P. (2017). Machine learning techniques for predicting crop photosynthetic capacity from leaf reflectance spectra. Molecular plant, 10(6), 878– 890.

17. Ji, F., Li, F., Hao, D., Shiklomanov, A. N., Yang, X., Townsend, P. A., Dashti, H., Nakaji, T., Kovach, K. R., Liu, H., et al. (2024). Unveiling the transferability of plsr models for leaf trait estimation: Lessons from a comprehensive analysis with a novel global dataset. New Phytologist, 243(1), 111–131.

18. Kaur, S., Kakani, V. G., Carver, B., Jarquin, D., & Singh, A. (2024). Hyperspectral imaging combined with machine learning for high-throughput phenotyping in winter wheat. The Plant Phenome Journal, 7(1), e20111.

19. Kole, C., Muthamilarasan, M., Henry, R., Edwards, D., Sharma, R., Abberton, M., Batley, J., Bentley, A., Blakeney, M., Bryant, J., et al. (2015). Application of genomics-assisted breeding for generation of climate resilient crops: Progress and prospects. Frontiers in plant science, 6, 563.

20. Kuhn & Max. (2008). Building predictive models in r using the caret package. Journal of Statistical Software, 28(5), 1–26. https://www.jstatsoft.org/index.php/jss/article/ view/v028i05

21. Kumar, L., Schmidt, K., Dury, S., & Skidmore, A. (2001). Imaging spectrometry and vegetation science. Imaging spectrometry: basic principles and prospective applications, 111–155.

22. Langridge, P., Braun, H., Hulke, B., Ober, E., & Prasanna, B. (2021). Breeding crops for climate resilience. Theoretical and Applied Genetics, 134(6), 1607–1611.

23. Meacham-Hensold, K., Montes, C. M., Wu, J., Guan, K., Fu, P., Ainsworth, E. A., Pederson, T., Moore, C. E., Brown, K. L., Raines, C., et al. (2019). High-throughput field phenotyping using hyperspectral reflectance and partial least squares regression (plsr) reveals genetic modifications to photosynthetic capacity. Remote Sensing of Environment, 231, 111176.

24. Meerdink, S. K., Roberts, D. A., King, J. Y., Roth, K. L., Dennison, P. E., Amaral, C. H., & Hook, S. J. (2016). Linking seasonal foliar traits to vswir-tir spectroscopy across california ecosystems. Remote Sensing of Environment, 186, 322–338.

25. Mevik, B.-H., & Wehrens, R. (2023). The pls package: Principal component and partial least squares regression in r. Journal of Statistical Software, 18(2), 1–24. 10.18637/jss.v018.i02

26. Rehman, T. U., Ma, D., Wang, L., Zhang, L., & Jin, J. (2020). Predictive spectral analysis using an end-to-end deep model from hyperspectral images for high-throughput plant phenotyping. Computers and Electronics in Agriculture, 177, 105713.

27. Serbin, S. P., Dillaway, D. N., Kruger, E. L., & Townsend, P. A. (2012). Leaf optical properties reflect variation in photosynthetic metabolism and its sensitivity to temperature. Journal of Experimental Botany, 63(1), 489–502.

28. Serbin, S. P., Singh, A., McNeil, B. E., Kingdon, C. C., & Townsend, P. A. (2014). Spectroscopic determination of leaf morphological and biochemical traits for northern temperate and boreal tree species. Ecological Applications, 24(7), 1651–1669.

29. Silva-Perez, V., Molero, G., Serbin, S. P., Condon, A. G., Reynolds, M. P., Furbank, R. T., & Evans, J. R. (2018). Hyperspectral reflectance as a tool to measure biochemical and physiological traits in wheat. Journal of Experimental Botany, 69(3), 483–496.

30. Tibshirani, R., & Friedman, J. (2001). The elements of statistical learning: Data mining, inference, and prediction. Springer.

31. Von Caemmerer, S. (2000). Biochemical models of leaf photosynthesis. Csiro publishing.

32. Wang, S., Guan, K., Wang, Z., Ainsworth, E. A., Zheng, T., Townsend, P. A., Li, K., Moller, C., Wu, G., & Jiang, C. (2021). Unique contributions of chlorophyll and nitrogen to predict crop photosynthetic capacity from leaf spectroscopy. Journal of experimental botany, 72(2), 341–354.

33. Wold, S., Sjöström, M., & Eriksson, L. (2001). Pls-regression: A basic tool of chemometrics. Chemometrics and intelligent laboratory systems, 58(2), 109–130.

34. Yan, Z., Guo, Z., Serbin, S. P., Song, G., Zhao, Y., Chen, Y., Wu, S., Wang, J., Wang, X., Li, J., et al. (2021). Spectroscopy outperforms leaf trait relationships for predicting photosynthetic capacity across different forest types. New Phytologist, 232(1), 134–147.

35. Yendrek, C. R., Tomaz, T., Montes, C. M., Cao, Y., Morse, A. M., Brown, P. J., McIntyre, L. M., Leakey, A. D., & Ainsworth, E. A. (2017). High-throughput phenotyping of maize leaf physiological and biochemical traits using hyperspectral reflectance. Plant physiology, 173(1), 614–626.

36. Yu, S., Fan, J., Lu, X., Wen, W., Shao, S., Guo, X., & Zhao, C. (2022). Hyperspectral techniqu combined with deep learning algorithm for prediction of phenotyping traits in lettuce. Frontiers in plant science, 13, 927832.

